# Guide RNA library representation results in gene essentiality prediction bias in genome-wide CRISPR screens

**DOI:** 10.1101/2025.09.07.673082

**Authors:** Paul Metz, Sofia Alves-Vasconcelos, Richard Wallbank, Joey Riepsaame, A. Bassim Hassan

**Author notes:** Corresponding author: Prof A B Hassan, Oxford Molecular Pathology Institute, Sir William Dunn School of Pathology, University of Oxford, South Parks Road, Oxford, OX1 3RE, UK. Department of Neurosurgery, Leiden University Medical Centre, Leiden, The Netherlands (Paul Metz).

## Abstract

Genome wide CRISPR-based perturbation screens are powerful discovery tools enabling the identification of novel gene dependencies through either gain or loss of function. While genome wide guide RNA (gRNA) libraries have advantages when using enAsCas12a, such as multiplex single gRNAs per gene, they may be subject to similar confounding factors that can affect the interpretation of large genome-wide datasets. Here, we examine the impact of these variables in over twenty enASCas12a multiple gRNA based perturbation screens performed using Humagne C, Humagne D and Inzolia libraries in human cells. We demonstrate that the choice of CRISPR library is often the most significant factor that influences genetic perturbation results, outweighing other variables such as either target cell lines or culture media conditions. A major contributor to this effect is gRNA representation bias within a given CRISPR library, where lower gRNA representation can lead to variable and more pronounced gene effect scores using either log fold change or Chronos analysis. These effects may be mitigated by using either multiple gRNA constructs per gene, by optimisation of CRISPR library production processes or by targeting with multiple independent gRNA libraries. Importantly, we also consider gRNA representation bias during CRISPR screen hit prioritisation. CRISPR library gRNA representation bias remains a major challenge in the interpretation of gene essentiality in perturbation screens.

## Introduction

CRISPR (Clustered Regularly Interspaced Short Palindromic Repeats) screening technologies have revolutionised the identification of gene dependent cellular vulnerabilities with unprecedented efficiency. For example, such screens have unveiled novel mechanisms underlying cancer pathology and highlighted promising therapeutic targets^1,2^. One significant technical advance has been the application of enhanced versions of the AsCas12a nuclease (enAsCas12a) coupled with multiplexed CRISPR libraries, enabling the delivery of multiple guide RNAs (gRNAs) per construct targeting either the same or different genes ^3, 4^. This approach facilitates targeting of two or more genomic loci simultaneously within a single cell, significantly increasing the scalability and efficiency of target disruption with a reduced cell number requirement.

Whilst CRISPR perturbation screens aim to provide clear discrimination between true dependencies (hits) and background noise, the reality is often far more complex ^5^. Extrinsic conditions can alter the range of gene dependencies, in particular the media that cells are cultured in ^6^. This has been highlighted recently with respect to essential gene and metabolic dependencies in CRISPR screens when comparing standard cell culture media (DMEM) with supplemented media that is physiologically closer to that observed in human serum ^7^. True target selection is not only dependent on such extrinsic conditions, but also on the intrinsic conditions within the screen. Hit prioritization remains largely non-standardized, as high ‘noise’ levels present significant challenges in distinguishing true positives and negatives from false positive and negatives. Single gRNAs with variable on-target efficiency have frequently been identified as a major contributor to this noise, with one study demonstrating that selective prioritisation of gRNAs with higher efficacy improves the specificity of true hit detection ^8^. Another large-scale screen analysis further highlighted additional noise sources, including variability in gRNA efficacy and batch effects, such as differences in CRISPR screen duration ^9^. Developments in gRNA design with the aid of tools such as CRISPpick, CHOPCHOP, and CRISPOR, have improved gRNA targeting efficiency through reducing off-target effects ^10, 11, 12^. Moreover, enhanced data analysis platforms that incorporate target copy number variation and time dependent assessment of gRNA representation, as demonstrated by CERES and Chronos, have all contributed to improved screen performance ^5, 13^. Despite these advances, there remains a significant degree of intrinsic and extrinsic noise that complicates the selection of true essentiality signals in CRISPR perturbation screens at lower scale. Here, we perform multiple screens with enAsCas12a libraries in human cells and investigate two key contributors to such noise: the choice of CRISPR library and the initial gRNA representation.

## Results

### Integrating variables impacting CRISPR screen

To begin evaluating the impact of different variables on CRISPR screen performance, we first developed a comprehensive workflow that incorporates two distinct human genome-wide enAsCas12a CRISPR libraries (Humagne C and D) in two different culture media (DMEM and HPLM) across four neural crest-derived cancer cell lines (Fig. 1a). Humagne C and D are separate genome-wide libraries (20,355 constructs per library) with two different guides per construct targeting a single gene. Having introduced enAsCas12a into human cell lines STS26T, ST88-14, 90-8 and iHSc (see Methods) using lentiviral transduction, we selected for bulk integration using Enhanced Green Fluorescent Protein (EGFP) based Fluorescence-activated cell sorting (FACS). We next tested media conditions by comparing culture in DMEM with human plasma like media (HPLM), which is considered to be more physiological ^6^. We observed cell growth rates to be slower in HPLM media such that we had to control for this observation in subsequent screens (Supplementary Figure 1). To mitigate potential silencing of the lentiviral inserted enAsCas12a expression following CRISPR library transduction in HPLM, we delayed the switch from DMEM to HPLM medium until after puromycin selection post transduction. In the comparison of DMEM and HPLM conditions, the early reference time point (t=0) was therefore set post-puromycin selection, rather than at the pDNA representation stage. To ensure a fair comparison across conditions, cells in each screen were allowed to proliferate to approximately 14 population doublings, accounting for differences in growth rates between cell lines and the effects of the culture medium. A total of 20 screens were performed and analysed in all cell lines using different cell line conditions, with at least one Humagne C and one Humagne D screen per cell line and condition. Guide abundance post-screen selection was determined with a minimum of three timepoints per screen across at least 14 population doublings.

**Figure 1.**
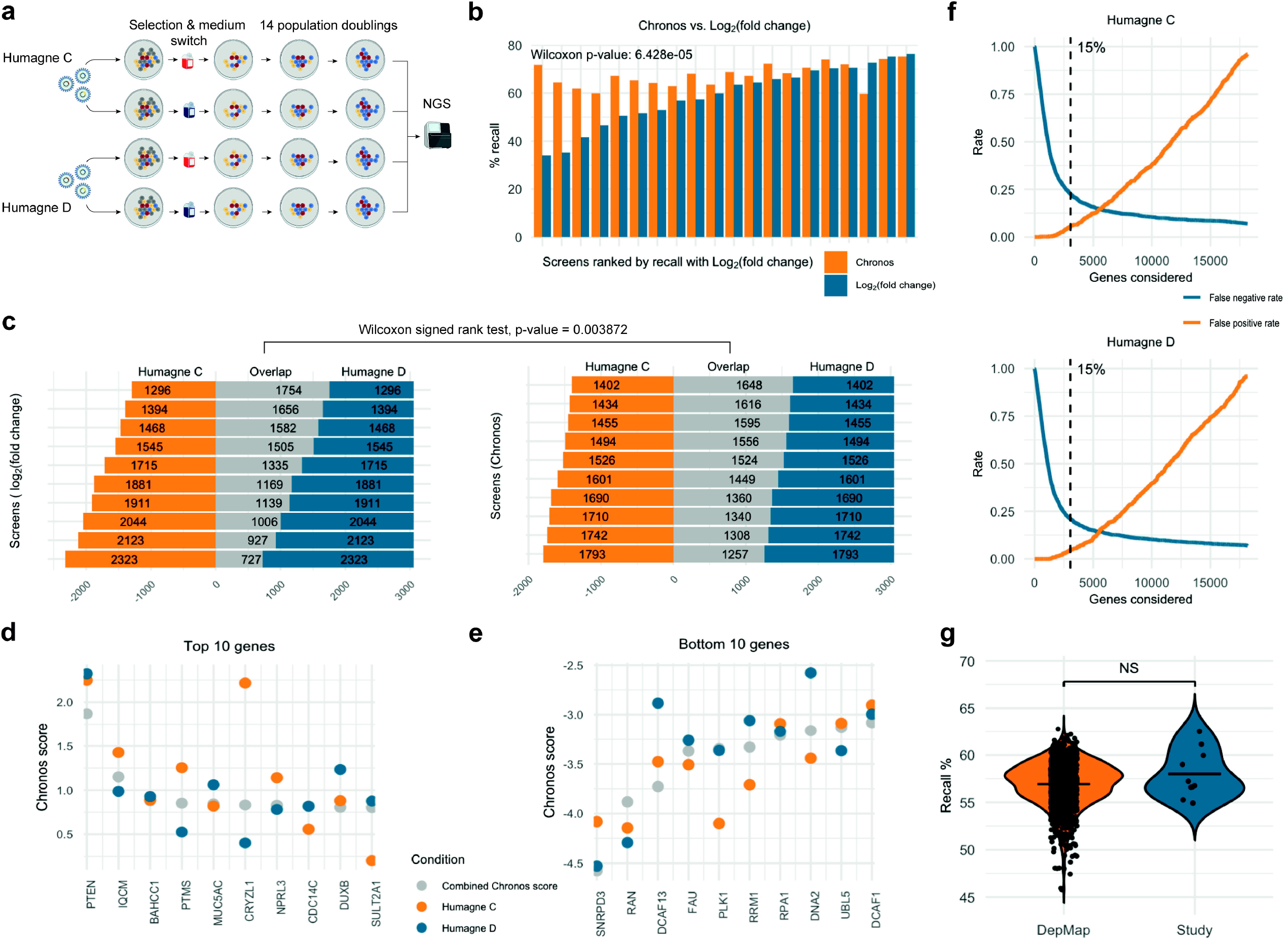
enAsCas12a CRISPR screen workflows and quality control. **a**. enAsCas12a CRISPR screen workflow using libraries Humagne C and D, and two different culture media, DMEM and HPLM. **b**. Bar graph comparing common essential gene recall of 20 CRISPR screens in this study following either log_2_(fold change) (blue) or Chronos analysis (orange). Log_2_(fold change) time points were selected based on best recall per time point while Chronos includes all available time points. Screens were ranked based on the recall obtained using log_2_(fold change). **c**. Horizontal stacked bar chart displaying overlapping and non-overlapping essential genes (bottom 15% genes) between CRISPR screens performed with Humagne C and Humagne D, analysed using the log_2_(fold change) or Chronos method. Screens are ranked by number of overlaps. Wilcoxon signed-rank test was used to compare the distribution of overlap counts between the log_2_(fold change) and Chronos groups. **d, e**. Dot plot of average Chronos scores per gene from Humagne C (orange), Humagne D (blue) and combined Humagne CD (grey). Genes displayed are the top 10 (**d**) and bottom 10 genes (**e**) based on Chronos score. **f**. Line plots displaying the false negative rate of recalled common essential genes (blue) and the false positive rate of negative control gRNAs (orange) calculated per number of bottom genes considered for Humagne C and Humagne D libraries. Chronos scores were averaged per gene across screens. **g**. Violin plots of recall percentages per CRISPR screen indicated as black dots, in both the DepMap dataset (orange violin) or this study (blue violin). Humagne C and D were analysed as a combined data set.

Various analysis tools are available for CRISPR screen data, with Chronos being the most recent and arguably the most comprehensive model to date ^13^. To benchmark Chronos against the conventional log_2_(fold-change) method, we compared the recall of common essential genes across screens using both approaches (Fig. 1b). Our analysis revealed that Chronos outperformed the log_2_(fold change) method in the majority of cases (17/20, 85 %), showing a significantly higher recall of common essential genes (p = 6.428E-05, paired Wilcoxon). Screens performed with the 26T cell linehad poorer recall, which we hypothesise may be partly due to reduced overall enAsCas12a expression in this cell line (Supplementary Fig. 2). Furthermore, we compared essential genes— defined as the bottom 15% of gRNA scores—across screens conducted under identical conditions, differing only in the CRISPR library used (Fig. 1c). This 15% threshold is based on the estimation that any given cell line has approximately this many true dependencies^13^. Notably, analysis with Chronos showed significantly greater similarity between these screens compared to the log_2_(fold change) method (Fig. 1c). Based on these findings, we proceeded with analyses of subsequent experiments were analysed using Chronos.

The CRISPR screens using Humagne C and D libraries can technically be analysed as a single experiment, with Chronos adjusting for batch effects across individual screens and generating a single gene score derived from multiple experiments. This approach can reduce the occurrence of false positives, as the combined gene score typically lies between the scores from experiments analysed separately (observed 73 % of the time, based on data from this study). This had the effect of often diminishing the extremity of the score—whether positive or negative (Fig. 1d and e). However, this strategy would be better extrapolated if false positives were observed more frequently than false negatives. Based on these data, however, we find that false negatives are 4.1-4.7 times more likely than false positives when considering the top and bottom 15% genes, for Humagne C and D, respectively (Fig. 1f). These data suggest that analysing the individual experiments as a unified screen may obscure true signals and result in the loss of meaningful biological hits.

To benchmark the quality of CRISPR screens in this study, common essential gene recall ratios were also compared to those of CRISPR screens presented in the DepMap dataset. No significant differences were detected between these two datasets (Fig. 1g).

### Effect of culture medium and CRISPR library on screen outcomes

Before assessing whether culture media alters the perturbation outcomes of Humagne C and D libraries, we evaluated the expression levels following RNA-sequencing of individual genes in cells cultured in HPLM versus DMEM for 21 days and revealed specific transcriptomic changes between the two media conditions (Fig. 2a). Across cell lines, 121 genes were differentially expressed, with 100 genes overexpressed and 21 under expressed in HPLM compared to DMEM, based on a log_2_(fold change) threshold of 1 and -1, respectively. The relatively modest number of differentially expressed genes suggest that culturing cells in HPLM versus DMEM induces significant but limited transcriptomic alterations common across cell lines.

**Figure 2.**
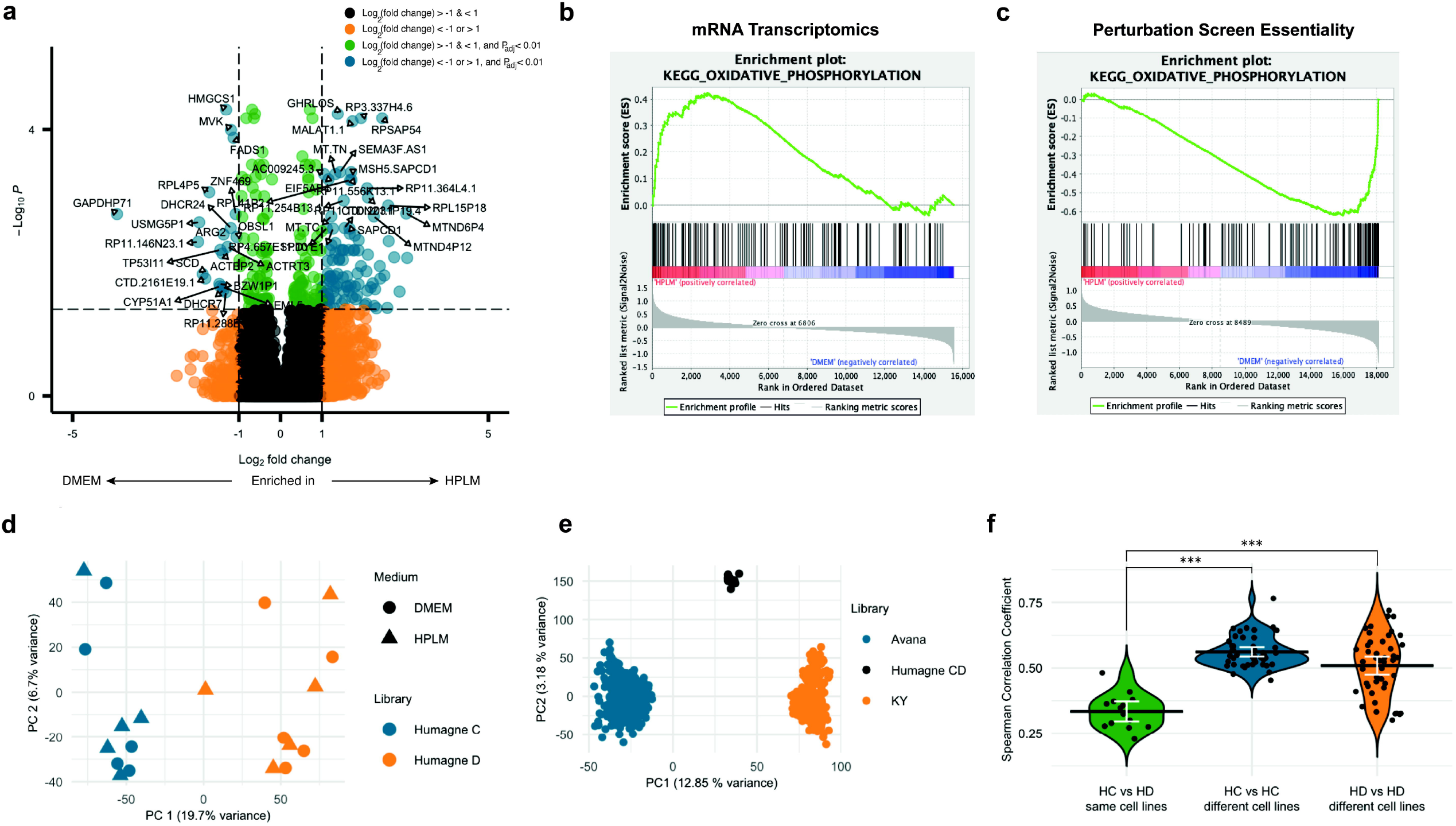
CRISPR library choice is major determinant of screen outcome. **a**. Volcano plot displaying genome-wide mRNA transcriptomics across target cell lines STS26T, 90-8, ST-8814 and iHSC, with two biological replicates each, grown in HPLM versus DMEM combined, with level and significance colour coded as indicated. Top differentially expressed genes are labelled by name with limited overlaps allowed. **b**. Enrichment plot of KEGG Oxidative Phosphorylation following GSEA of combined mRNA transcriptomics of STS26T, 90-8, ST-8814 and iHSC cells cultured in HPLM versus DMEM. The complete list of significantly enriched sets in either medium is shown in Supplementary Table 1. **c**. GSEA plots of significantly (FDR q value (<25%)) differential essentiality genes with Hallmark and KeggLecacy gene sets, where a positive correlation reflects relatively higher Chronos scores for genes across HPLM versus DMEM CRISPR screens in this set. The complete list of significantly enriched sets in either medium is shown in Supplementary Table 2. **d**. Dot plot displaying principal component analysis (PCA) on all CRISPR library screens performed (N = 20) in this study separately analysed for Humagne C (blue) and D (orange) libraries. The variance explained by principal components is indicated on the side of each axis. PC1: principal component 1, PC2: principal component 2. **e**. Dot plot displaying principal component analysis (PCA) on all CRISPR library screens from DepMap, with Humagne C and D analysed by Chronos as if one experiment as the data is provided in this way, with dots colour coded according to library as indicated. PC1: principal component 1, PC2: principal component 2. **f**. Violin plots displaying Spearman’s Rho (R) values of correlations between the same cell lines screened with Humagne C and Humagne D (green) compared to all possible correlations between different cell lines and conditions screened with the same library, Humagne C (blue) or Humagne D (red). Means per group are displayed by a horizontal black line and mean confidence intervals by white brackets. Wilcoxon rank-sum tests were performed to compare the distribution of Spearman’s Rho (R) values between groups. Asterisks indicate significance: ^*^ p < 0.05, ^**^ p < 0.01, ^***^ p < 0.001, NS not significant.

Some of the selected transcriptomic changes are reflected in the enrichment of specific gene expression sets in either HPLM or DMEM, as identified by experimentally curated Gene Set Enrichment Analysis (GSEA, Supplementary Table 1). Notably, metabolic reprogramming is evident through various gene set enrichment in both culture media. For instance, GESA in HPLM showed enrichment in mRNA transcriptomics gene sets associated with oxidative phosphorylation (Fig. 2b) and respiratory electron transport (Supplementary Table 1), indicating enhanced mitochondrial activity and energy production. The elevated levels of oxidative phosphorylation in HPLM may be due to its lower glucose concentration (5 mM) compared to the 25 mM in high-glucose DMEM, the standard medium for these cell lines. The reduced glucose availability in HPLM may shift cellular metabolism toward a greater dependence on mitochondrial energy production^14^.

To determine whether CRISPR screens could uncover dependencies associated with these gene expression gene sets, GSEA was then performed on the results of the CRISPR perturbation screen essentiality data from cell lines cultured in DMEM and HPLM. For these analysis, data from Humagne C and D screens were combined to enhance statistical power. In this context, enrichment indicates relatively higher gene essentiality or a reduced growth effect, while depletion suggests the opposite. Interestingly, these analyses reveal that the vast majority of differentially essential gene sets do not overlap with differentially expressed transcriptomics gene sets from the non-screen analyses (Supplementary Table 1 and 2). This suggests that differential essentiality is more likely to occur in genes of which the expression is tightly regulated. However, some screen essential gene sets do correlate with differentially expressed genes. For instance, in line with the previous transcriptomic analysis, genes related to oxidative phosphorylation were more essential in HPLM, aligning with the idea that cells may depend more on mitochondrial energy production when glucose levels are lower (Fig. 2c) ^14^. Importantly, there were no instances where higher gene set essentiality was associated with lower expression within the same medium.

Several other variables are believed to play a role in determining quality and outcomes of CRISPR screen experiments, including the specific target cell line, duration of the screen and CRISPR library choice. We next performed principal component analysis (PCA) on data from Humagne C and Humagne D screens from this study (Fig. 2d). This revealed that principal component 1 (PC1) effectively distinguishes screens based primarily on library choice, rather than either the cell line model or culture medium used, while explaining 19.7% of the total variance. To determine whether this observation was specific to either the methodology employed in this study or relevant to a broader context, we conducted a similar PCA on the CRISPR screen data available from DepMap. Interestingly, we observed a comparable clustering pattern across CRISPR datasets generated using the Avana, KY, and HCD libraries (Fig. 2e). Notably, the overlap of cell lines between the Avana and KY screens (204 in total) provided evidence that the choice of CRISPR library is a more significant determinant of screen outcomes than the target cell line itself. To further test this hypothesis, we calculated Pearson correlation rho values between screens performed on the same target cell lines but using different CRISPR libraries (Humagne C vs. Humagne D) and compared these to correlations between screens using the same CRISPR libraries (e.g. Humagne C vs. Humagne C) but across different target cell lines. All possible comparisons were included, except those involving the same cell line (Fig. 2f). Although some individual cases may show stronger correlations when screens are performed with the same cell line rather than the same library, the majority of the comparisons indicate that screens performed with the same CRISPR library exhibit significantly higher correlations, supporting the possibility that library choice may have a greater impact in determining essentiality outcome compared to the cancer cell line model.

### Examining CRISPR library representation bias

Another key difference between CRISPR libraries is the variation in gRNA construct representation, referred to here as plasmid DNA (pDNA) representation. Drawing from the law of large numbers ^15^, that states that a larger number of samples tested converges to the hypothetical true mean of the distribution, we hypothesised that the differences in representation of plasmids targeting the same gene across different libraries may also contribute to the differential perturbation outcomes observed between different CRISPR libraries.

To evaluate the impact of pDNA representation on CRISPR screen outcomes, we first examined the raw data by displaying the relationship between pDNA count and average log_2_(fold change) across Humagne C and D CRISPR library screens, separately (Fig. 3a and 3b). The visualisation of this relationship suggests that pDNAs with lower representation in the library tend to exhibit greater variation with either more extreme positive or negative scores compared to those with higher representation which show a narrower distribution of scores. Notably, bins containing pDNAs with lower representation show larger standard deviations, confirming more variability in gene effect scores, whereas bins with higher representation exhibit smaller standard deviations, reflecting less variability. As visualisation of raw data may simply mask more frequent pDNA representation towards the lower end of the counts, mean log_2_(fold change) values per gene across screens were placed in equally sized bins and the medians per bin plotted in order to remove this potential bias (Fig. 3c and 3d). This analysis showed that lower pDNA representation is indeed associated with more dramatic positive or negative log_2_(fold change) gene effect scores.

**Figure 3.**
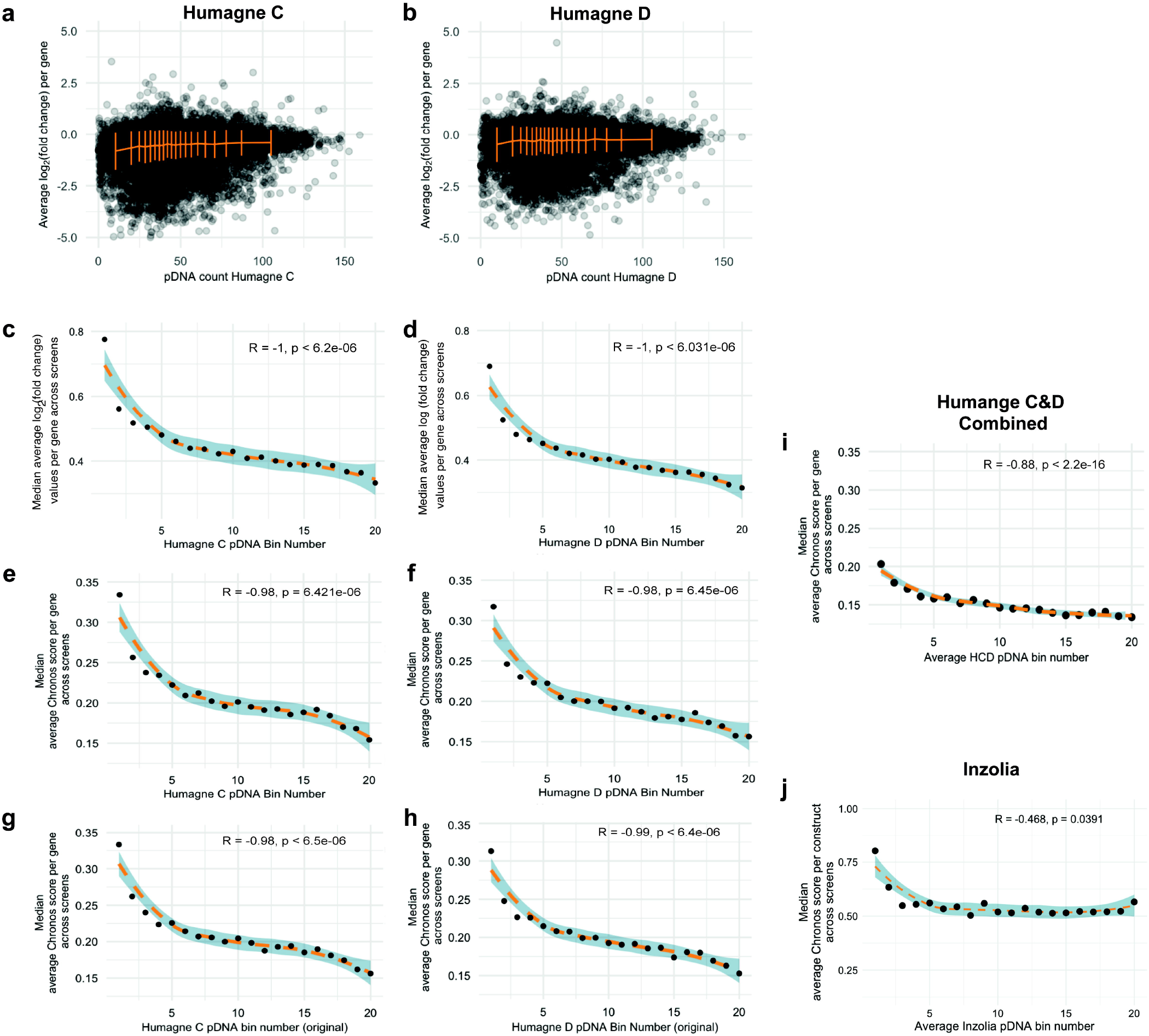
CRISPR library representation presents source of bias. **a**. Dot plots displaying the relationship between the pDNA count and the average log2(fold change) per gene across Humagne C screens. Standard deviations (orange) are calculated across 20 equally sized bins containing average log2(fold change) values per gene across screens. **b**. Same as (a.) but for Humagne D screens from this study. **c**. Dot plot and locally estimated scatterplot smoothing (LOESS) (orange dashed line) with 95% confidence interval (blue) of the relationship between ranked Humagne C pDNA count bins and the median of the mean absolute log2(fold change) values per gRNA across Humagne C screens, per pDNA count bin. **d**. Same as (c.), but for Humagne D screens from this study. **e**. Dot plot and locally estimated scatterplot smoothing (LOESS) (orange dashed line) with 95% confidence interval (blue) of the relationship between ranked Humagne C pDNA count bins and the median of the mean absolute Chronos scores per gene across Humagne C screens, per pDNA count bin. A Spearman’s rank correlation test was performed to test for significance, with Rho (R) and p value displayed on the plot. **f**. Same as for (e.) but for Humagne D screens from this study. **g**. Based on the same analysis as (e.), but in relation to original pDNA count before library amplification. **h**. Same as (g.), but for Humagne D screens from this study. **i**. Dot plot and LOESS (orange dashed line) with 95% confidence interval of the relationship between pDNA count bins, based on the average pDNA count per gene across Humagne C and D, and the median of the mean absolute Chronos score per gene across Humagne CD (analysed by Chronos as if one library) screens, per pDNA count bin. A Spearman’s rank correlation test was performed to test for significance, with Rho (R) and p value displayed on the plot. **j**. Same as for (i) but for the Inzolia screens.

To evaluate whether Chronos indirectly corrects for the effect of pDNA representation and to further quantify this relationship, we next calculated the average absolute Chronos scores per gene across CRISPR screens. Here, average Chronos scores per gene were placed in equally sized bins based on their associated pDNA representation. We then performed Pearson correlation analyses on the median Chronos scores per gene for each bin (Fig. 3e and 3f). For both Humagne C and D libraries, the correlations were highly significant, indicating that Chronos does not (effectively) adjust for the variability associated with pDNA representation.

To determine whether this pDNA representation effect is a result of either in-house CRISPR library amplification or inherent to the original, unamplified library, we binned the average Chronos scores per gene across screens according to the pDNA representation of the original CRISPR library (Fig. 3g and 3h). Notably, these results closely mirrored those obtained using the pDNA counts from the amplified library, suggesting that the observed effect is intrinsic to the original library and not merely an artefact of the amplification process. Furthermore, the highly significant correlation between the original pDNA and post-amplification library representation suggests that library amplification, if performed correctly, should barely influence the impact of representation bias (Supplementary Figure 3).

As previously discussed, Chronos can also analyse Humagne C and D screens as if they were performed as a single experiment. We hypothesised that as the two gRNA cassettes in each of the Humagne C and D libraries are amplified separately, the effect of low pDNA representation in one library could potentially either mitigate or counterbalance a normal representation in the other. In order to evaluate this, Chronos determined single scores per gene from both equivalent Humagne C and D screens. Then, average scores per gene were calculated across different CRISPR screens and binned according to their associated average pDNA representation (average of Humagne C and Humagne D pDNA count). Again, Pearson correlation analysis was used to evaluate the relationship between the median score per pDNA bin and pDNA representation (Fig. 3i). Although this approach reduced the effect compared to analysing Humagne C and D separately, as measured by Spearman’s correlation, the overall relationship in terms of representation bias remained strong (Rho: -0.88) and highly significant (p-value < 2.2e-16). Thus, while Chronos may mitigate representation bias by generating a single gene effect score that combines data from two separate libraries, the influence of pDNA library representation bias is still evident.

To evaluate if the effect was also present in data sets obtained using different CRISPR libraries than Humagne C and D, we next performed two Inzolia-based CRISPR library screens (Supplementary Figure 4). The Inzolia sgRNA library is also enAsCas12a based with two constructs per gene containing the same four gRNAs but in a different sequence order ^4^. The library also contains paralog gene sgRNAs in the same construct facilitating multiplex gene targeting. Although less significant, a similar relationship was observed between Chronos scores and pDNA count in the Inzolia library screens (Fig. 3j). This observation may possibly be due to the lower number of screens performed and current lack of an Inzolia Chronos training dataset.

To assess further whether the observed pDNA representation effect is either specific to the Humagne C and D libraries or is a broader characteristic of CRISPR libraries, we analysed the DepMap dataset using a similar methodology. The Avana and KY libraries typically contain four and five gRNA constructs per gene, respectively, while Chronos still generates a single score per gene even if there are multiple gRNAs. To determine if the pDNA representation effect persists despite the presence of multiple gRNA constructs per gene, an aspect that could potentially strongly dilute the impact of outliers with low representation, we averaged the pDNA counts across constructs for each gene. The mean absolute Chronos scores for each gene across screens were then categorised into equally sized bins based on the averaged pDNA count. Then, median scores across genes were computed per bin. The analysis across the Avana, KY, and Humagne CD screens from the DepMap dataset consistently revealed the same effect, characterised by high Spearman’s correlation coefficients (Rho) and significant p-values of near 0 (Fig. 4a-f). As predicted, these findings indicate that the inclusion of multiple gRNA constructs per gene in a library does not completely attenuate the impact of the initial pDNA representation on CRISPR screen outcomes, and that this effect seems to impact all CRISPR libraries tested.

**Figure 4.**
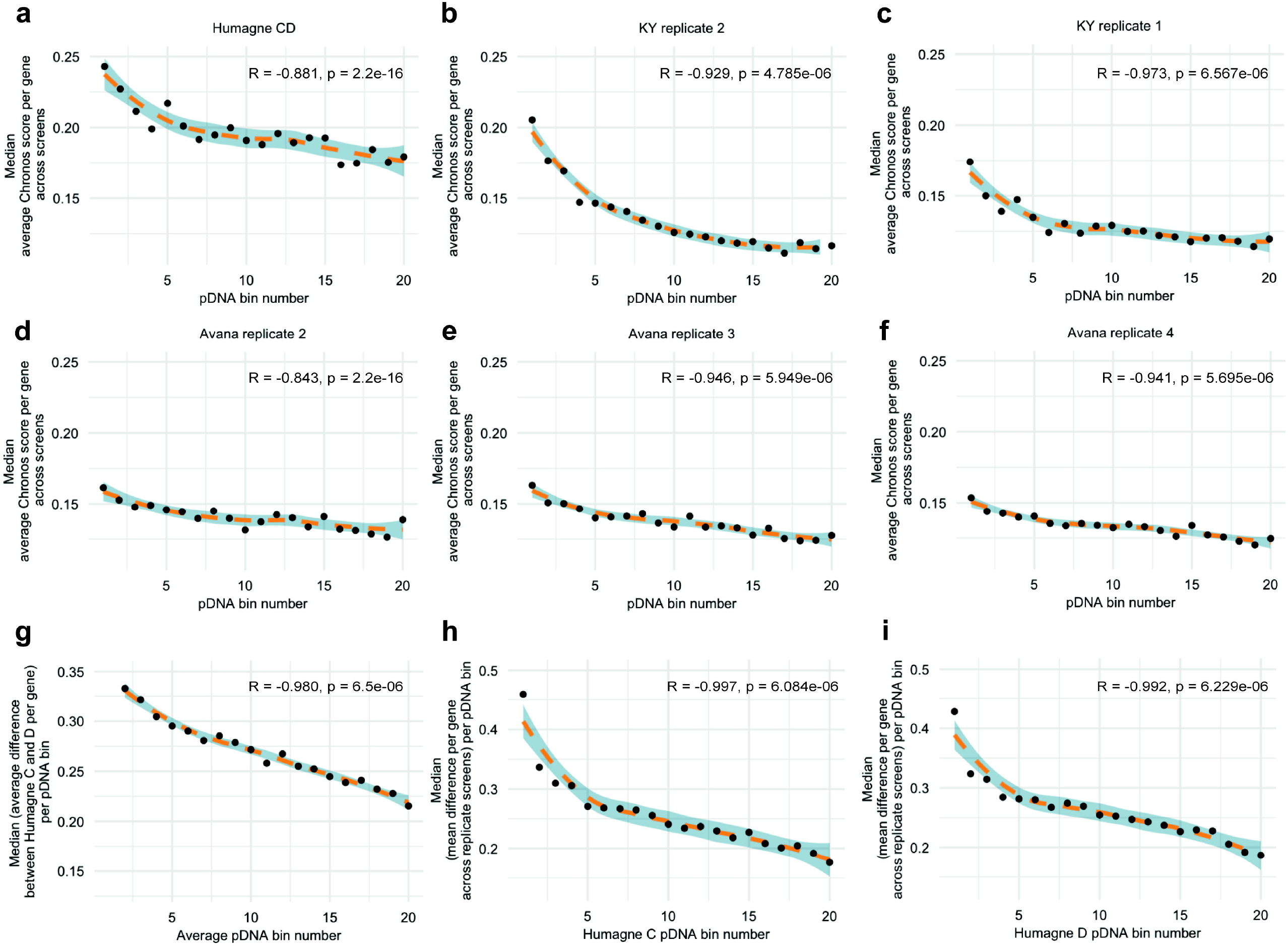
CRISPR library representation bias present in external datasets (DepMap). **a**. Dot plot and LOESS (orange dashed line) with 95% confidence interval of the relationship between average pDNA count bins and the median of the mean absolute Chronos scores per gene across DepMap’s Humagne CD screens, per pDNA count bin. A Spearman’s rank correlation test was performed to test for significance, with Rho (R) and p value displayed on the plot. **b-f**. Same as for (a.) but for DepMap’s KY batch 2 (b.), KY batch 3 (c.), Avana batch 1 (d.), Avana batch 2 (e.), Avana batch 3 (f.). **g**. Dot plot and LOESS (orange dashed line) with 95% confidence interval of the relationship between average pDNA count bins and the median of the mean absolute difference between Humagne C and D Chronos scores per gene across screens. A Spearman’s rank correlation test was performed to test for significance, with Rho (R) and p value displayed on the plot. **h**. Dot plot and LOESS (orange dashed line) with 95% confidence interval of the relationship between Humagne C pDNA count bins and the median of the mean absolute difference between first and second replicate Chronos scores per gene across replicate Humagne C screens, per pDNA count bin. A Spearman’s rank correlation test was performed to test for significance, with Rho (R) and p value displayed on the plot. **i**. Same as in (h.) but for Humagne D CRISPR screens.

To investigate the relationship between pDNA representation and the previously observed differences between CRISPR libraries, we next calculated the absolute differences in Chronos scores between Humagne C and D libraries across various cell line models. These differences were averaged per gene across multiple screens, converted to absolute values, and subsequently binned according to associated average Humagne C and D pDNA counts. A Spearman’s correlation analysis of the median values per bin revealed a very strong and highly significant correlation, confirming that pDNA representation significantly influences the variability in outcomes when different CRISPR libraries are applied (Fig. 4g). Furthermore, differences observed between replicate screens (utilising the same library) were similarly attributable to the pDNA effect to a comparable degree (Fig. 4h and 4i).

### False positives prediction based on library representation bias

In an attempt to further quantify the pDNA representation effect, we predicted the number of false positive essential genes depending on the pDNA representation threshold below which associated outcomes may be deemed relatively unreliable. First, the 15% lowest scoring genes, referred to as essential genes, were selected based on Chronos scores from Humagne C and D screens with iHSC cells cultured in DMEM (Fig. 5a, b, respectively). We then calculated the ratio of genes that were deemed essential compared to all genes below each possible pDNA threshold (Fig 5c, d). The number of predicted false positives was calculated as follows. We first determined both the expected number of essential genes below each pDNA threshold (15% of this subset of genes) and the observed number of essential genes (as part of the general 15% most essential genes) below the pDNA threshold (Fig. 5e, f). The predicted false positives were the subtraction of the latter from the former. Dependent on the threshold of pDNA representation, different numbers of false positives are estimated. For example, when setting the pDNA count threshold at 25, roughly 100 genes of that subset are estimated to be false positive for the screens depicted (Fig. 5e, f). Additionally, by binning genes based on pDNA count in equal bins, one can calculate the number of false positive and negative essential genes per bin and so estimate the total number of false positive and false negative genes in total (Fig. 5g, h). Based on the analysis displayed, an estimated 187 (6.9%) and 143 (5.3%) out of 2703 essential genes (15%) are false positive and the same number of genes is falsely classified as negative, totalling a number of 374 and 287 genes for these Humagne C and D screens, respectively (Fig. 5g, h). We made similar observations for all other CRISPR screens performed with Humagne C and D and Inzolia this study (Supplementary Figure 5-13).

**Figure 5.**
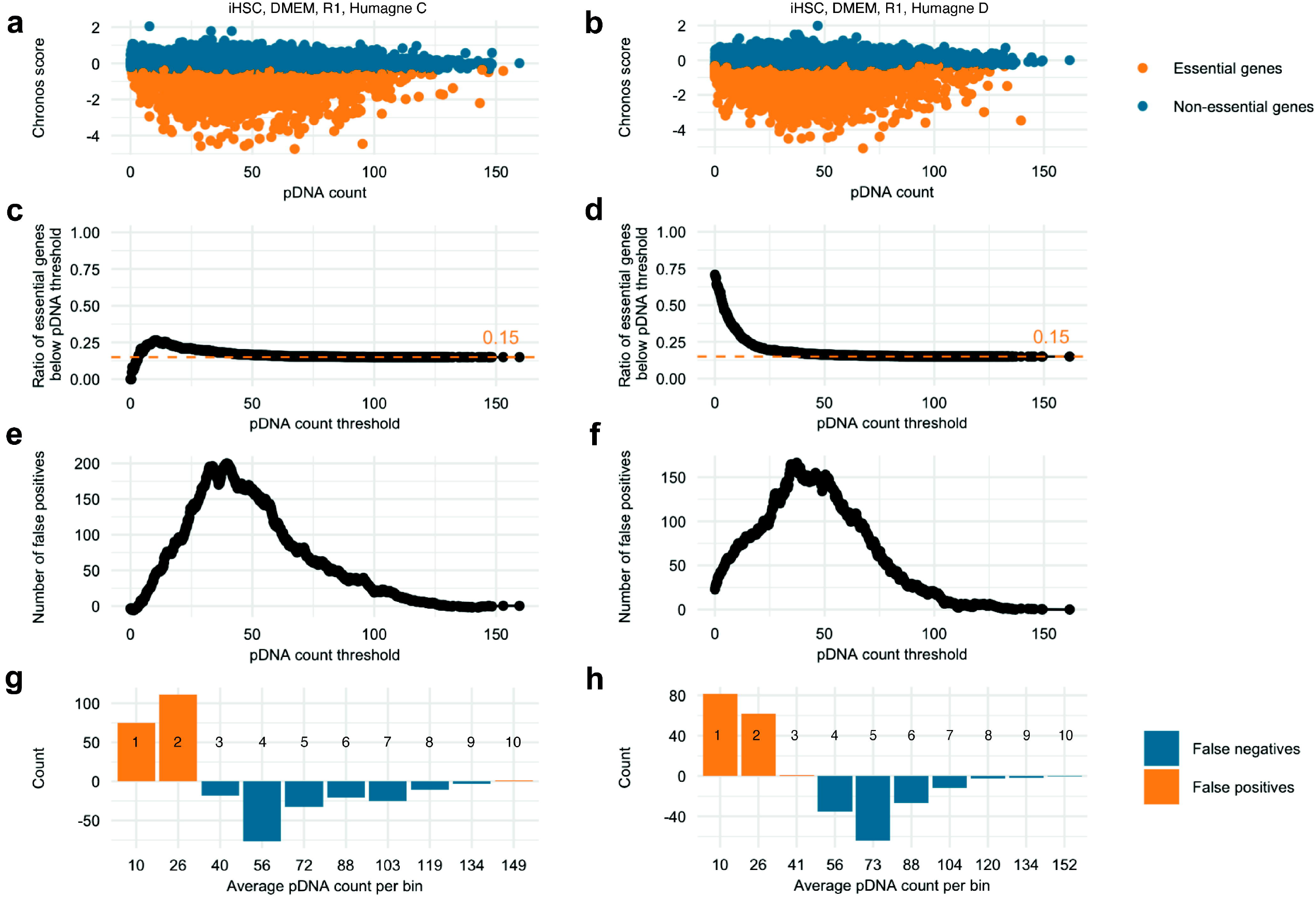
Estimation of false positive and negative results coming from pDNA representation bias. **a**. Dot plot of the relationship between Chronos scores and pDNA counts in iHSC, DMEM, replicate 1, Humagne C CRISPR screen. The bottom 15% genes based on Chronos score are highlighted in orange. **b**. Same as in (a.) but for the equivalent Humagne D screen. **c**. Dot plot depicting the ratio of found essential genes and the total number of genes below each possible pDNA threshold for the iHSC, DMEM, replicate 1, Humagne C CRISPR screen. The dashed orange line indicates the predicted 15% essential genes threshold if essential genes were equally distributed across pDNA representation **d**. Same as in c but for the equivalent Humagne D screen. **e**. Dot plot showing the predicted number of false positives below each possible pDNA threshold for the iHSC, DMEM, replicate 1, Humagne C CRISPR screen. **f**. Same as in (e.) but for the equivalent Humagne D screen. **g**. Bar graph of the estimated number of false positive and negative essential genes per pDNA bin for the iHSC, DMEM, replicate 1, Humagne C CRISPR screen, with the x axis showing the average number of pDNA counts associated with each bin and the number on each bar indicating the bin number. **h**. same as (g.) but for the equivalent Humagne D screen.

### Reducing library representation bias

A recent study demonstrated that the difference in pDNA representation within CRISPR libraries may be reduced by optimising key steps in library production, such as oligo synthesis direction, the number of PCR cycles, and annealing temperature (Tm) ^16^. The primary aim of the work was to improve the scalability of CRISPR screens, given their labour-intensive nature in terms of cell numbers, yet these optimisations may also enhance pDNA representation and mitigate bias in screens. To investigate these findings, we compared the impact of pDNA representation on gene effect scores in a CRISPRi screen using libraries optimised for representation from the above-mentioned study ^16^, with data from the Weisman group’s original, ‘non-optimised’ libraries ^17^. By analysing both datasets, we sought to determine whether the optimised library development workflow reduced the impact of pDNA representation on gene effect scores. As observed in our study and corroborated by external data from DepMap, the correlation between pDNA representation and gene effect scores from the non-optimized workflow from the Weisman group data was close to 1 (Rho replicate 1 = -0.98 and Rho replicate 2 = -0.96), and highly significant (Fig. 5a and 5b). In contrast, the correlation between pDNA representation and gene effect scores was significantly lower in the optimised library workflow, with the average Pearson correlation dropping from -0.97 (N = 2) to -0.76 (N = 3) (Fig. 5c-f). Notably, because pDNA representation counts were not available for both the PCR optimised and non-optimised libraries, we used gRNA counts measured after puromycin selection as a proxy for pDNA representation and as an early reference time point. Therefore, although significant, these data may not show the complete effect of an optimised PCR library production workflow on the pDNA representation bias effect.

Lastly, we attempted to perform a post-experimental correction for the pDNA representation bias by calculating z-scores for gene effect scores within each pDNA bin. While this approach yielded significant improvements in showing similarity between replicate CRISPR screens and Humagne C versus Humagne D screens, the effect appeared to be only marginal (Fig. 6g, h). No significant improvement was shown in common essential gene recall (Fig 6i).

**Figure 6.**
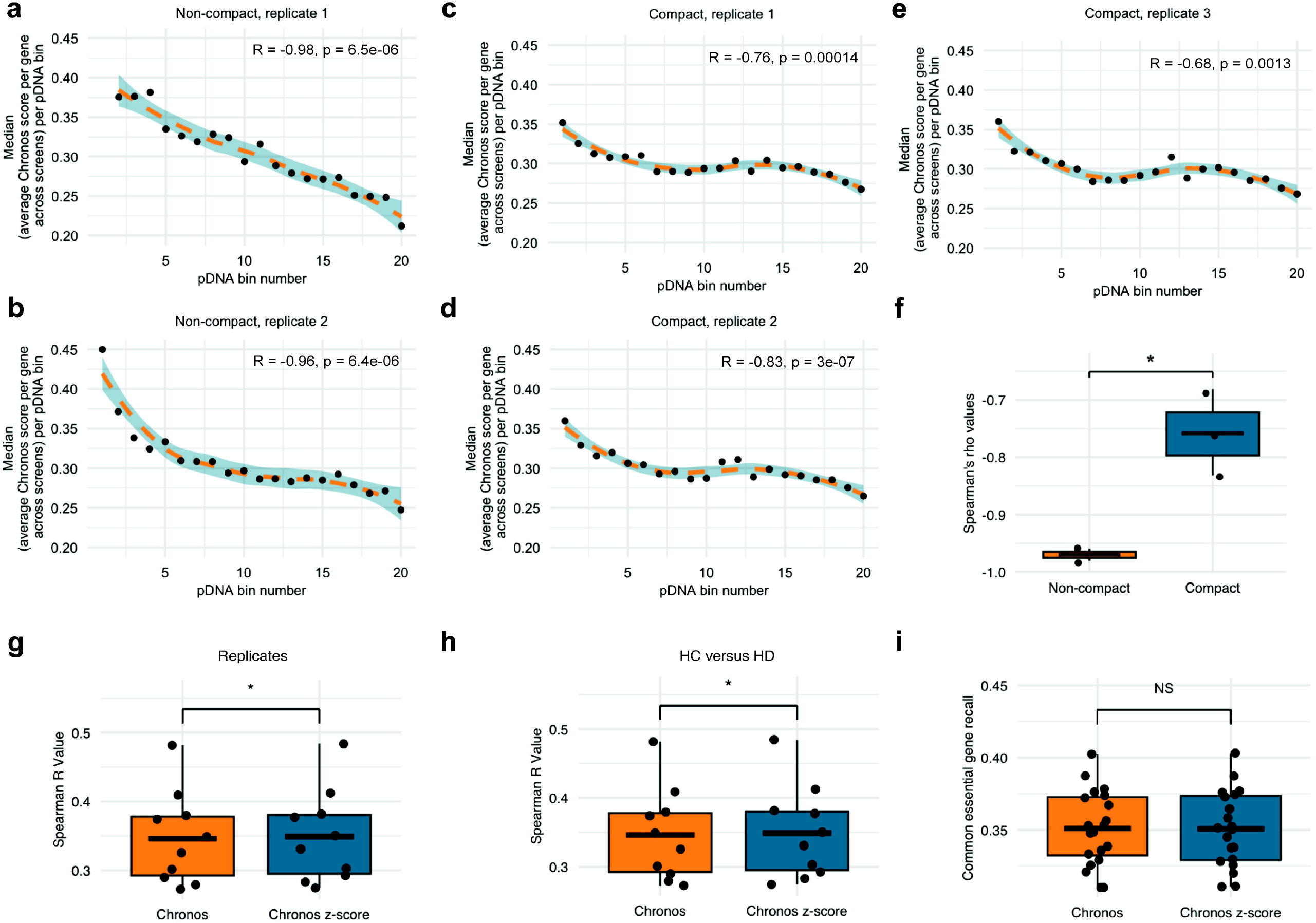
CRISPR library production optimisation reduces representation bias. **a**. Dot plot and LOESS (orange dashed line) with 95% confidence interval of the relationship between the median of the mean absolute Chronos scores per gene for replicate 1 from the CRISPR screen performed with a library generated following the ‘non-optimised’ workflow^17^. **b**. Same as (a.) but for replicate 2. **c.-e**. Same as a., but data from CRISPR screens performed with libraries that were generated following the optimised workflow^16^. **f**. Box plots of the Spearman’s Rho correlations from (a-e) comparing the ‘non-optimised’ to the ‘optimised’ library production workflow. **g**. Box plot displaying Spearman’s Rho (R) values of correlations between replicate screens analysed using Chronos (orange) regularly or Chronos after which z-scores are calculated per gene per pDNA bin (blue). **h**. Same as (g.) but for the comparison between Humagne C and D equivalent screens. **i**. Box plot displaying common essential recall rate values of all Humagne C and D screens performed in this study, analysed using Chronos (orange) regularly or Chronos after which z-scores are calculated per gene per pDNA bin (blue). **f-i**. Means are represented by horizontal black lines. A Wilcoxon signed-rank test was performed to compare the values between groups. Asterisks indicate significance: ^*^ p < 0.05, ^**^ p < 0.01, ^***^ p < 0.001, NS means not significant.

## Discussion

In the present study, we have analysed data from twenty CRISPR screens performed in our own laboratory across four human cell lines, in two types of culture medium, using three distinct CRISPR libraries, Humagne C, Humagne D, and an additional 2 screens using the Inzolia library in a further cell line. The aim of the analysis was to identify extrinsic and intrinsic variables that could impact on screen results. Our screen data are of similar quality to CRISPR screen data available from DepMap based on the recall of common essential genes, and so are comparable. Similarly, we confirmed that Chronos analysis significantly improves the quality of most CRISPR screen data compared to Log_2_ (fold change) and was utilised to uniformly analyse all screen data ^13^.

We observed that extrinsic conditions, such as the use of different medium types DMEM and HPLM, led to differential dependencies, but that these dependencies did not always correlate with differential gene expression. One of the few GSEA gene sets that was found to be both overexpressed in HPLM and more essential in HPLM was that of oxidative phosphorylation (KEGG gene set), which may suggest a higher dependency of cells on mitochondrial energy production due to lower glucose levels in HPLM versus DMEM. Overall, the effects of culture medium on cellular dependencies were found to be confined to discrete metabolic gene sets based on our data.

In contrast, we observed that the choice of genome-wide CRISPR library was a major factor that influenced CRISPR screen essentiality outcomes. The results from CRISPR screens performed with Humagne C and D libraries can be perfectly separated during a PCA analysis, as demonstrated by in-house and external datasets (DepMap). Furthermore, CRISPR library choice also appeared of greater importance in determining outcomes than extrinsic factors like culture medium conditions, as demonstrated with DMEM and HPLM media in this study, and, perhaps more importantly, the cell line model.

Our analysis identified that gRNA representation in the CRISPR library as a significant intrinsic factor influencing CRISPR screen results. Specifically, lower gRNA representation results in a wider variability in either positive or negative gRNA representation changes during the screen, leading to skewed stronger gene effect scores during data analysis, with both conventional log_2_ (fold change) as well as Chronos. Conversely, higher gRNA representation was associated with less dramatic gene effect scoring. The gRNA representation bias was present across all screens performed in this study, including the novel all-in-one enAsCas12a Inzolia library, as well as in external datasets (DepMap), spanning three different CRISPR libraries (Avana, KY and Humagne CD). These data suggest that gRNA representation bias seems to play a major role in differences observed between Humagne C and D CRISPR library essentiality results, where lower gRNA representation is associated with larger variation between results from Humagne C and D screens, compared to gRNAs that have a higher initial representation. This observation also needs to be more strongly considered in addition to other factors that impact differences between CRISPR libraries proposed by other studies, including differences in gRNA efficiencies and batch effects^9^.

To quantify the effect of the gRNA representation bias, we estimated the number of false positives and negative essential genes based on the expected number of essential genes and the observed number of essential genes in different gRNA representation categories. In line with our findings, the number of essential genes, as defined by the bottom 15% scoring genes is higher amongst genes associated with gRNAs that have a relatively low initial representation. By estimating the number of false positive and negative essential genes per gRNA representation bin, we predicted that the total number of wrongly classified (false positive or negative) genes in screen data from this study varies between 6 and 23%.

Post-experimental correction of this effect proved difficult. Calculating z-scores from Chronos scores based on their associated gRNA representation category only resulted in marginal improvement of similarity between results from replicates and screens performed with different libraries. The recall of common essential genes was not improved following this correction. Different approaches should be tested to see if there are ways to retrospectively improve the data and screen coverage by correcting for gRNA presentation bias, although it is unlikely that the corrections will fully compensate for what is a structural and inherent feature of library design and cell number dependency. Aside from PCR cycle optimisation and PCR primer sequence bias, reducing gRNA representation bias would either include the use of a greater cell number to compensate for low gRNA representation or to utilise independently generated guide libraries, that fundamentally differ in pDNA representation, combined with eliminating the low pDNA bins from the analysis. For a given screen coverage and cell number, these would still have cost-efficiency implications and result in either multiple library screens per experiment or even pooled but separately generated gRNA libraries with high cell numbers.

Based on the findings in these studies, we also propose that hit selection from CRISPR screen data should include the consideration of gRNA representation in the library, as guides with a lower pDNA result in more extreme gene effect scores and can lead to the selection of false positive candidate genes. At the very least, we suggest that where hit genes with similar gene effect scores are associated with different gRNA representation, hit genes with the highest gRNA representation should be prioritised for downstream validation as they are less likely to reflect an effect coming from gRNA representation bias. Additionally, as our data predicts that a substantial proportion of genes are false positives, one may consider disregarding genes of which the initial gRNA representation is below a certain threshold. Based on data in this study for example, a normalised pDNA count threshold of 25 could be considered as an arbitrary cut-off, but this empirical suggestion is likely to be context based.

## Supporting information

Cover Letter

## Acknowledgements

We thank Stuart Brown for statistical discussion and members of the Dunn School for technical advice.

## Author contributions

PM co-designed and performed all Humagne C and D screen experiments, completed informatics and statistical analysis and wrote the manuscript, SAV performed and provided Inzolia library data, RW and JR provided technical advice, and ABH conceived of the project, obtained grant funding and wrote the manuscript with PM and SAV.

## Materials and Methods

### Mammalian cell culture

The mammalian cell lines used in this study are listed in Supplementary Table 3 and authenticated following short tandem repeat (STR) profiling (Supplementary Table 4). MPNST cell lines STS26T, ST88-14 and 90-8 were kindly provided by Professor Eduard Serra (Germans Trias i Pujol Research Institute, Barcelona, Spain), the immortalised Human Schwann Cell line (iHSC; ipn02.3 1-λ) by Professor Nancy Ratner (Cincinnati Children’s Hospital Medical Center, Cincinnati, USA), and LentiX Hek293T cell line by Professor Carvalho (University of Oxford, UK). Specifically, the iHSC line used in this study was made from the same population of Schwann cells as ipn02.3 2λ, developed by the group of Professor Wallace (University of Florida), but transduced with 1 instead of 2 µL of each virus (retro and lentivirus) carrying the hTERT and mCdk4 immortalisation genes ^18^. The osteosarcoma cell line HOS (derived from a 13 year-old female caucasian) was acquired from Culture Collections (ECACC 87070202) and the osteosarcoma cell line OHS (derived from a 13 year old male) was gifted by collaborator Professor Leonardo Meza-Zepeda, from the Radium Hospital (NRH), University of Oslo. Cell lines tested negative for mycoplasma contamination by polymerase chain reaction (PCR) using primers 5’-YGCCTGVGTAGTAYRYWCGC-3’ and 5’-GCGGTGTGTACAARMCCCGA-3’^19^. MPNST and iHSC cells were cultured in DMEM complemented with 10% FBS and Penicillin/Streptomycin (100 Units/mL) and passaged by first washing with Dulbecco’s phosphate-buffered saline (DPBS) (D8573-500 mL Sigma Life Sciences) and then treatment with TrypLE Express (CAT #: 12604-013, ThermoFisher Scientific) when reaching a confluency of 70-90%. The HOS cell line was cultured in Eagle’s Minimum Essential Medium (EMEM) supplemented with 2mM Glutamine, 1% Non-Essential Amino Acids (NEAA) and 10% Foetal Bovine Serum (FBS). The OHS cell line was cultured in RPMI1640 supplemented with 10% Foetal Calf Serum (FCS) and 1x glutamax. Counting cells was performed with AO-DAPI Solution 13 following manufacturer’s instructions. To quantify population doublings, a standard curve was established correlating percentage of cell growth to population doublings, where the initial time point is defined as 100% at 0 population doublings. Each subsequent doubling increases the population count by a factor of two, reflecting a linear progression of 1 population doubling per doubling in percentage. After each passage, the percentage increase in cell count relative to the previous time point was measured and converted into population doublings using this standard curve.

### CRISPR library amplification

CRISPR libraries Humagne C and D, and Inzolia, were diluted to achieve a concentration of 40 ng/µL in a final volume of 50 µL. For each electroporation, 5 µL (200 ng) of the diluted CRISPR pDNA library was added to 35 µL of MegaX competent cells. Electroporation was performed in 0.1 cm cuvettes at 2.0 kV, 200 ohms, and 25 µF. After electroporation, 465 µL of SOC medium was immediately added, and the mixture was transferred to a round-bottom tube containing 2.5 mL of prewarmed SOC medium. This process was repeated three times for each library. The cells were then incubated in SOC medium at 37°C with shaking for 1 hour. Following recovery, the cultured cells were combined into a sterile tube, gently mixed, and a small sample was diluted and plated to estimate colony counts. The remaining recovered culture was added to 500 mL of 2XYT + AMP and incubated overnight at 37°C in a shaking incubator. On the next day, colonies were counted, with 333 colonies for Library C and 402 colonies for Library D, both above the generally recommended threshold of 100 colonies per construct proceeding with library DNA purification^20^. Isolation and purification of the amplified CRISPR libraries was performed by using a Qiagen Plasmid Maxi Kit (Qiagen: 12162) on the cultured bacteria.

### CRISPR library validation

The sgRNA region of Humagne C and D was amplified using primers designed for sequencing with 41 bp cycles, requiring one of the sgRNA pairs to be within 41 bp of the Illumina adapter. Primers with known annealing sites (P5_ARGON and P7_KERMIT) were used, incorporating a stagger region for library diversity and an index for sample barcoding. The sgRNA region of Inzolia was amplified with primers P5_ARGON and P7_MISSPIGGY. The sequences for the forward and reverse primers used in this process are provided in Supplementary Table 5. Following PCR (Supplementary Table 6), DNA products were analysed using gel electrophoresis on a 2% agarose gel to confirm the presence of the expected 350 bp bands. The samples were then quantified using Qubit for Illumina sequencing at 500X coverage using a NextSeq sequencing platform at Azenta Life Science ^21^. The validated libraries were then used for lentiviral production and subsequent CRISPR screens.

### Lentivirus Production via PEI-Mediated Transfection

LentiX HEK293FT cells were cultured to 70-80% confluence in T175 flasks and transfected with a plasmid mixture containing 2.64 µg of pMD2.G, 5.28 µg of psPAX2, and 10.56 µg of the lentiviral transfer plasmid in 506.9 µL of serum-free DMEM. The plasmid solution was combined with 151.7 µL of PEI solution, vortexed, and incubated at room temperature for 10 minutes to allow for complex formation. The transfection mixture was then added to the cells in 25 mL of DMEM supplemented with 10% FBS. After 24 hours, the medium was replaced with fresh DMEM containing 10% FBS. Forty-eight hours post-transfection, the lentiviral supernatant was harvested by filtering twice, using 0.45 μm PES filters, followed by snap-freezing and storage at -80 °C.

### Generation of enAsCas12a-stable cell lines transduction

To generate enAsCas12a-stable cell lines, EGFP was inserted into pRDA_174 via an auto-cleaving peptide sequence T2A in the same connected to enAsCas12a in the same reading frame via regular cloning methods. Lentivirus was produced as described above. Target cells were seeded and transduced in suspension to achieve a transduction efficiency of approximately 30%. Cells were first expanded and then sorted for EGFP positives based on fluorescence-activated cell sorting (FACS).

### Genome-wide CRISPR knockout screens

This workflow largely follows a previously described genome-wide CRISPR screen workflow described by Zhang’s Group (Eli and Edythe L. Broad Institute of MIT and Harvard, USA)^20^. Cryopreserved cells were thawed and expanded until they reached a volume of 200-300 × 10^6 cells. Cell density and viability were assessed using acridine orange and DAPI (AO-DAPI) staining. Cells were then seeded at a minimum density of 6.12 × 10^6^ cells per T175 flask, with the number of flasks adjusted to ensure a post-selection yield of at least 500 cells per construct within the CRISPR library (> 10,500 E6 cells), in 25 mL of DMEM supplemented with 10% FBS (antibiotic-free). To optimise lentiviral transduction, 25 µL of a 10 mg/mL polybrene solution was added to each flask, resulting in a final polybrene concentration of 10 µg/mL. A predetermined volume of 160 µL of Humagne C or D lentivirus was then added to each flask to achieve a transduction efficiency of approximately 30%. After 24 hours of incubation, the transduction medium was removed and replaced with fresh standard medium, containing puromycin at a concentration optimised for the specific target cell line (Supplementary Table 7). The cells were incubated for an additional 72 hours. After these 72 hours, and each subsequent passage, the cells were detached, counted, and the population doublings (PDs) were calculated. A multiple of 12 × 10^6^ cells were pelleted, washed with ice cold PBS and stored at - 80 °C for later genomic DNA (gDNA) extraction, ensuring full coverage of over 500 cells per CRISPR library construct per pellet (> 10.5 × 10^6^ cells). Of the remainder, a total of 12 × 10^6^ cells (> 500x coverage) were then reseeded across 9 x T175 flasks in fresh DMEM supplemented with 10% FBS. This process was repeated at least every four days until the cells reached approximately 14 population doublings, at which point a final harvest was conducted, and the screen was terminated. For CRISPR screens in HPLM, a similar workflow was utilised, except that at the first passage, right after puromycin selection, medium was switched from standard medium to HPLM complemented with 10% FBS and 100 Units/mL Penicillin and Streptomycin. HPLM medium was replaced with every passage as with DMEM medium. Genomic DNA was isolated from up to 12×10^6^ cultured cells using the Mag-Bind® Blood & Tissue DNA HDQ 96 Kit (Omega Bio-Tek). PCR amplification of the gRNA regions was performed using the NEBNext® Ultra™ II Q5® Master Mix (NEB), optimised for high-fidelity amplification. The PCR master mix for each sample included 10 µM concentrations of both forward and reverse primers, which were designed to be compatible with next-generation sequencing (NGS) workflows, allowing for efficient pooling (Supplementary Table 8). The master mix also included 1X NEB Q5 High-Fidelity Master Mix and nuclease-free water. All available gDNA per sample was used, with a maximum of 20 ng of gDNA per 100 µL PCR reaction volume. The total reaction volume was a multiple of 100 µL, based on the amount of gDNA available for each sample. These reactions underwent a touchdown PCR programme (Supplementary Table 9). Post-PCR, samples were run on a 1% agarose gel and appropriate bands isolated using a Zymoclean Gel DNA Recovery Kit (Zymo Research). Samples were quantified using a Qubit™ dsDNA Quantification Assay Kit (Invitrogen™, Catalogue number: Q32851) and sequenced at 500 reads per library construct (> 10.5×10^6^ reads per sample) on an Illumina NextSeq platform using 150 base pair paired-end reads.

### CRISPR screen data analysis using Log_2_(fold change) and Chronos

For Log_2_ (fold change) analysis, FASTQ files were first analysed using FASTQC and subsequently processed with the MAGeCK count function as part of the MAGeCK RRA algorithm^22^, aligning sequencing reads to a reference gRNA library to generate a count matrix reflecting gRNA abundance across different time points with the library gRNA representation as the first time point reference. The counts per gRNA were normalised using Reads Per Million (RPM). Subsequently, log_2_ (fold change) values and downstream analyses were performed in R ^23^. Chronos was performed following the developer’s instructions with the library gRNA representation as the first time point reference ^24^. The ‘nan_outgrowths’ function in Chronos was applied to correct for gRNAs exhibiting excessive outgrowth^13^. Chronos was performed on separate or combined Humagne C and D experiments in a pretrained mode, with the model configured using parameters derived from training on the DepMap data from Humagne C and D CRISPR screens. Essential genes were identified by ranking genes and selecting those within the bottom 15%, both for log_2_(fold change) and Chronos scores. The overlap between these selected genes and the list of common essential genes from the Trevor Hart group, was then calculated to determine recall ^25^. To evaluate the quality of the CRISPR screens conducted in this study, they were compared to DepMap CRISPR screens (Gene effect scores, Supplementary Table 10) based on recall of common essential genes. DepMap also provided a list of common non-essential genes (Common non-essential Genes, Supplementary Table 10). The effect of gRNA representation in DepMap datasets was determined using the pDNA counts from the DepMap gRNA count matrix (DepMap gRNA Count Matrix, Supplementary Table 10).

### RNA sequencing analysis

RNA sequencing was performed on cell lines STS26T, 90-8, ST-8814, and iHSC, sequenced at Azenta Life Science ^21^. Quality control of the raw sequencing data was conducted using FastQC ^26^. Adapter removal and trimming of low-quality reads were performed with Trimmomatic (v0.39) ^27^. The cleaned reads were then aligned to the reference genome (Homo sapiens GRCh38) using STAR (version 2.7.8)^28,29^. For gene quantification, FeatureCounts (part of SUBREAD version 2.0.3) ^29^ was employed to generate raw count matrices. Differential gene expression analysis was carried out using the DESeq2 workflow, following the guidelines outlined in the DESeq2 vignette available at Bioconductor ^30^. Gene set enrichment analysis (GSEA) was performed utilising Signal2Noise metrics for HPLM versus DMEM conditions ^31^. GSEA focussed on Hallmark, KEGG Legacy, and Reactome gene sets to assess functional enrichment of differentially expressed genes ^31^.

