## Supplementary Tables for "Guide RNA library representation results in gene essentiality prediction bias in genome-wide CRISPR screens"

**Supplementary Table 1. GSEA of combined transcriptomics of STS26T, 90-8, ST-8814 and iHSC cells cultured in HPLM versus DMEM.**

| Gene set | Enriched in | FDR q value (<25%) |
|---|---|---|
| Hallmark_cholesterol_homeostasis | DMEM | 0.024 |
| Hallmark_peroxisome | DMEM | 0.072 |
| Hallmark_wnt_beta_catenin_signaling | DMEM | 0.245 |
| Hallmark_androgen_response | DMEM | 0.214 |
| Hallmark_adipogenesis | DMEM | 0.194 |
| Hallmark_mtorc1_signaling | DMEM | 0.191 |
| Kegg_ribosome | HPLM | 0.14 |
| Kegg_protein_export | HPLM | 0.225 |
| Kegg_hematopoietic_cell_lineage | HPLM | 0.179 |
| Kegg_type_i_diabetes_mellitus | HPLM | 0.181 |
| Kegg_parkinsons_disease | HPLM | 0.155 |
| Kegg_oxidative_phosphorylation | HPLM | 0.155 |
| Kegg_biosynthesis_of_unsaturated_fatty_acids | DMEM | 0.055 |
| Kegg_ppar_signaling_pathway | DMEM | 0.078 |
| Kegg_endometrial_cancer | DMEM | 0.136 |
| Kegg_inositol_phosphate_metabolism | DMEM | 0.102 |
| Kegg_mtor_signaling_pathway | DMEM | 0.121 |
| Kegg_citrate_cycle_tca_cycle | DMEM | 0.103 |
| Kegg_notch_signaling_pathway | DMEM | 0.105 |
| Kegg_aminoacyl_trna_biosynthesis | DMEM | 0.111 |
| Kegg_acute_myeloid_leukemia | DMEM | 0.105 |
| Kegg_steroid_biosynthesis | DMEM | 0.115 |

| | | |
|---|---|---|
| Kegg_glycosaminoglycan_biosynthesis_heparan_sulfate | DMEM | 0.15 |
| Kegg_phosphatidylinositol_signaling_system | DMEM | 0.141 |
| Kegg_non_small_cell_lung_cancer | DMEM | 0.131 |
| Kegg_b_cell_receptor_signaling_pathway | DMEM | 0.228 |
| Kegg_fc_gamma_r_mediated_phagocytosis | DMEM | 0.212 |
| Kegg_lysine_degradation | DMEM | 0.209 |
| Kegg_vegf_signaling_pathway | DMEM | 0.208 |
| Kegg_peroxisome | DMEM | 0.205 |
| Reactome_sars_cov_1_modulates_host_translation_machinery | HPLM | 0.156 |
| Reactome_srp_dependent_cotranslational_protein_targeting_to_membrane | HPLM | 0.139 |
| Reactome_eukaryotic_translation_elongation | HPLM | 0.191 |
| Reactome_activation_of_the_mrna_upon_binding_of_the_cap_binding_complex_and_eifs_and_subsequent_binding_to_43s | HPLM | 0.183 |
| Reactome_eukaryotic_translation_initiation | HPLM | 0.249 |
| Reactome_respiratory_electron_transport | HPLM | 0.222 |
| Reactome_response_of_eif2ak4_gcn2_to_amino_acid_deficiency | HPLM | 0.204 |
| Reactome_regulation_of_expression_of_slits_and_robos | HPLM | 0.193 |
| Reactome_nonsense_mediated_decay_nmd | HPLM | 0.194 |
| Reactome_complex_i_biogenesis | HPLM | 0.206 |
| Reactome_influenza_infection | HPLM | 0.234 |
| Reactome_selenoamino_acid_metabolism | HPLM | 0.226 |
| Reactome_sars_cov_1_host_interactions | HPLM | 0.213 |
| Reactome_respiratory_electron_transport_atp_synthesis_by_chemiosmotic_coupling_and_heat_production_by_uncoupling_proteins | HPLM | 0.22 |

**Supplementary Table 2. GSEA of combined CRISPR perturbation screen data from STS26T, 90-8, ST-8814 and iHSC cells cultured in HPLM versus DMEM.**

| Gene set | Essential in | FDR q value (<25%) |
|---|---|---|
| HALLMARK_UNFOLDED_PROTEIN_RESPONSE | DMEM | 0.202 |
| HALLMARK_OXIDATIVE_PHOSPHORYLATION | HPLM | 0.002 |
| HALLMARK_FATTY_ACID_METABOLISM | HPLM | 0.095 |
| KEGG_PROTEIN_EXPORT | DMEM | 0.208 |
| KEGG_PARKINSONS_DISEASE | HPLM | 0.003 |
| KEGG_OXIDATIVE_PHOSPHORYLATION | HPLM | 0.001 |
| KEGG_ALZHEIMERS_DISEASE | HPLM | 0.004 |
| KEGG_HUNTINGTONS_DISEASE | HPLM | 0.004 |
| KEGG_CITRATE_CYCLE_TCA_CYCLE | HPLM | 0.059 |
| REACTOME_TRNA_PROCESSING_IN_THE_NUCLEUS | DMEM | 0.084 |
| REACTOME_COMPLEX_I_BIOGENESIS | HPLM | 0.013 |
| REACTOME_AEROBIC_RESPIRATION_AND_RESPIRATORY_ELECTRON_TRANSPORT | HPLM | 0.009 |
| REACTOME_RESPIRATORY_ELECTRON_TRANSPORT | HPLM | 0.007 |
| REACTOME_COMPLEX_IV_ASSEMBLY | HPLM | 0.195 |
| REACTOME_ACTIVATION_OF_ANTERIOR_HOX_GENE_IN_HINDBRAIN_DEVELOPMENT_DURING_EARLY_EMBRYOGENESIS | HPLM | 0.192 |
| REACTOME_METABOLISM_OF_PORPHYRINS | HPLM | 0.179 |
| REACTOME_FORMATION_OF_ATP_BY_CHEMIOSMOTIC_COUPLING | HPLM | 0.175 |
| REACTOME_PHASE_2_PLATEAU_PHASE | HPLM | 0.19 |
| REACTOME_GLYOXYLATE_METABOLISM_AND_GLYCINE_DEGRADATION | HPLM | 0.194 |
| REACTOME_INTERLEUKIN_7_SIGNALING | HPLM | 0.177 |
| REACTOME_OXIDATIVE_STRESS_INDUCED_SENESCENCE | HPLM | 0.226 |
| REACTOME_ERBB2_REGULATES_CELL_MOTILITY | HPLM | 0.233 |
| REACTOME_RECOGNITION_AND_ASSOCIATION_OF_DNA_GLYCOSYLASE_WITH_SITE_CONTAINING_AN_AFFECTED_PURINE | HPLM | 0.222 |
| REACTOME_MEIOTIC_RECOMBINATION | HPLM | 0.213 |
| REACTOME_BASE_EXCISION_REPAIR_AP_SITE_FORMATION | HPLM | 0.202 |
| REACTOME_INHIBITION_OF_DNA_RECOMBINATION_AT_TELOMERE | HPLM | 0.193 |
| REACTOME_CHROMATIN_MODIFICATIONS_DURING_THE_MATERNAL_TO_ZYGOTIC_TRANSITION_MZT | HPLM | 0.188 |
| REACTOME_MEIOSIS | HPLM | 0.178 |
| REACTOME_FORMATION_OF_THE_BETA_CATENIN_TCF_TRANSACTIVATING_COMPLEX | HPLM | 0.18 |

| | | |
|---|---|---|
| REACTOME_ASSEMBLY_OF_THE_ORC_COMPLEX_AT_THE_ORIGIN_OF_REPLICATION | HPLM | 0.173 |
| REACTOME_PRC2_METHYLATES_HISTONES_AND_DNA | HPLM | 0.173 |
| REACTOME_RUNX1_REGULATES_GENES_INVOLVED_IN_MEGAKARYOCYTE_DIFFERENTIATION_AND_PLATELET_FUNCTION | HPLM | 0.167 |
| REACTOME_DNA_METHYLATION | HPLM | 0.163 |
| REACTOME_CONDENSATION_OF_PROPHASE_CHROMOSOMES | HPLM | 0.162 |
| REACTOME_REGULATION_OF_TLR_BY_ENDOGENOUS_LIGAND | HPLM | 0.171 |
| REACTOME_BUDDING_AND_MATURATION_OF_HIV_VIRION | HPLM | 0.176 |
| REACTOME_MEIOTIC_SYNAPSIS | HPLM | 0.17 |
| REACTOME_ERCC6_CSB_AND_EHMT2_G9A_POSITIVELY_REGULATE_RRNA_EXPRESSION | HPLM | 0.176 |
| REACTOME_REGULATION_OF_ENDOGENOUS_RETROELEMENTS_BY_PIWI_INTERACTING_RNAS_PIRNAS | HPLM | 0.175 |
| REACTOME_BASE_EXCISION_REPAIR | HPLM | 0.19 |
| REACTOME_SENESCENCE_ASSOCIATED_SECRETORY_PHENOTYPE_SASP | HPLM | 0.191 |
| REACTOME_RMTS_METHYLATE_HISTONE_ARGININES | HPLM | 0.187 |
| REACTOME_REGULATION_OF_ENDOGENOUS_RETROELEMENTS_BY_THE_HUMAN_SILENCING_HUB_HUSH_COMPLEX | HPLM | 0.197 |
| REACTOME_TRANSCRIPTIONAL_REGULATION_OF_GRANULOPOIESIS | HPLM | 0.21 |
| REACTOME_FORMATION_OF_THE_NEPHRIC_DUCT | HPLM | 0.227 |
| REACTOME_DEFENSINS | HPLM | 0.227 |
| REACTOME_REGULATION_OF_INNATE_IMMUNE_RESPONSES_TO_CYTOSOLIC_DNA | HPLM | 0.229 |
| REACTOME_SURFACTANT_METABOLISM | HPLM | 0.224 |
| REACTOME_REGULATION_OF_ENDOGENOUS_RETROELEMENTS_BY_KRAB_ZFP_PROTEINS | HPLM | 0.223 |
| REACTOME_REGULATION_OF_PYRUVATE_DEHYDROGENASE_PDH_COMPLEX | HPLM | 0.23 |
| REACTOME_MATERNAL_TO_ZYGOTIC_ TRANSITION_MZT | HPLM | 0.235 |
| REACTOME_DISEASES_OF_PROGRAMMED_ CELL_DEATH | HPLM | 0.231 |
| REACTOME_CITRIC_ACID_CYCLE_TCA_CYCLE | HPLM | 0.245 |
| REACTOME_STIMULI_SENSING_CHANNELS | HPLM | 0.248 |

**Supplementary Table 3. Cell lines used in this study.**

| Cell line | Origin, Disease | Standard medium* | Reference |
|---|---|---|---|
| STS26T | Human melanoma, sporadic | DMEM + 10% FBS + Pen/Strep | [33] |
| ST-8814 | Human MPNST, NF1 | DMEM + 10% FBS + Pen/Strep | [34] |
| 90-8 | Human MPNST, NF1 | DMEM + 10% FBS + Pen/Strep | [35] |
| iHSC | Human, no disease | DMEM + 10% FBS + Pen/Strep | [36] |
| LentiX Hek293T | Human Embryonic Kidney | DMEM + 10% FBS + Pen/Strep | Takara Bio, CAT # 632180 |
| HOS | Human osteosarcoma | RPMI + 10% FBS + 1% GlutaMax | ATCC #CRL-1543 |
| OHS | Human osteosarcoma | RPMI + 10% FBS + 1% NEAA + 2mM GlutaMax | Obtained from Radium Hospital (NRH), University of Oslo |

**Supplementary Table 4. Short tandem repeat (STR) profiling of target cell lines**

| Marker | STS26T | 90-8 | ST-8814 | iHSC | LentiX Hek293T |
|---|---|---|---|---|---|
| AMEL | X,X | X,X | X,Y | X,X | X,X |
| CSF1PO | 10,13 | 12,12 | 9,12 | 10,11 | 11,12,13 |
| D13S317 | 9,10 | 8,8 | 12,12 | 12,13 | 12,13,14,15 |
| D16S539 | 12,13 | 9,9 | 13,13 | 12,12 | 9,14,13 |
| D18S51 | 17,18 | 14,18 | 12,12 | 13,16 | 17,18 |
| D19S433 | 14,14 | 13,16 | 13,14 | 14,14 | 17,18,19 |
| D21S11 | 30,31 | 30,30 | 29,32.2 | 30.2,32.2 | 28,29,30.2,31.2 |
| D2S1338 | 20,20 | 16,24 | 17,23 | 16,17 | 19,19 |
| D3S1358 | 14,14 | 15,15 | 15,18 | 17,18 | 15,16,17 |
| D5S818 | 11,12 | 11,11 | 12,13 | 8,11 | 8,9 |
| D7S820 | 8,11 | 11,11 | 8,8 | 8,12 | 11,11 |
| D8S1179 | 13,14 | 10,12 | 14,14 | 12,15 | 11,12,13,14,15 |
| FGA | 22,23 | 25,25 | 21,21 | 23,24 | 23,23 |
| TH01 | 6, 9.3 | 6,6 | 9,9 | 6,7 | 7,9.3 |
| TPOX | 8,8 | 11,11 | 11,12 | 8,12 | 11,11 |
| vWA | 17,17 | 17,17 | 16,16 | 15,17 | 16,17,18,19,20 |

**Supplementary Table 5. Primers used for CRISPR library validation**

| Primer Name | Sequence (5' to 3') |
|---|---|
| P5_ARGON_1 | AATGATACGGCGACCACCGAGATCTACACTCTTTCCCTACACGACGCTCTTCCGATCTTTGTGGAAAGGACGAAACACCG |
| P5_ARGON_2 | AATGATACGGCGACCACCGAGATCTACACTCTTTCCCTACACGACGCTCTTCCGATCTCTTGTGGAAAGGACGAAACACCG |
| P5_ARGON_3 | AATGATACGGCGACCACCGAGATCTACACTCTTTCCCTACACGACGCTCTTCCGATCTGCTTGTGGAAAGGACGAAACACCG |
| P5_ARGON_4 | AATGATACGGCGACCACCGAGATCTACACTCTTTCCCTACACGACGCTCTTCCGATCTAGCTTGTGGAAAGGACGAAACACCG |
| P7_KERMIT_1 | CAAGCAGAAGACGGCATACGAGATTCGCCTTAGTGACTGGAGTTCAGACGTGTGCTCTTCCGATCTTCTACTATTCTTTCCCCTGCACTGT |
| P7_KERMIT_2 | CAAGCAGAAGACGGCATACGAGATCTAGTACGGTGACTGGAGTTCAGACGTGTGCTCTTCCGATCTTCTACTATTCTTTCCCCTGCACTGT |
| P7_KERMIT_3 | CAAGCAGAAGACGGCATACGAGATGCTCAGGAGTGACTGGAGTTCAGACGTGTGCTCTTCCGATCTTCTACTATTCTTTCCCCTGCACTGT |
| P7_KERMIT_4 | CAAGCAGAAGACGGCATACGAGATAGGAGTCCGTGACTGGAGTTCAGACGTGTGCTCTTCCGATCTTCTACTATTCTTTCCCCTGCACTG |
| P7_MISSPIGGY_1 | CAAGCAGAAGACGGCATACGAGATATTACTCGGTGACTGGAGTTCAGACGTGTGCTCTTCCGATCTACCGACTCGGTGCCACTTTTTCAA |
| P7_MISSPIGGY_2 | CAAGCAGAAGACGGCATACGAGATTCCGGAGAGTGACTGGAGTTCAGACGTGTGCTCTTCCGATCTACCGACTCGGTGCCACTTTTTCAA |

**Supplementary Table 6. PCR programme used for CRISPR library validation**

| Stage | Temperature (°C) | Time | Number of Cycles |
|---|---|---|---|
| Initial Denaturation | 98 | 03:00 | 1x |
| Denaturation | 98 | 00:15 | 5x (Stage B) |
| Annealing | 65 | 00:30 | 5x (Stage B) |
| Extension | 72 | 00:30 | 5x (Stage B) |
| Denaturation | 98 | 00:15 | 8x (Stage C) |
| Annealing | 63 | 00:30 | 8x (Stage C) |
| Extension | 72 | 00:30 | 8x (Stage C) |
| Denaturation | 98 | 00:15 | 12x (Stage D) |
| Annealing | 61 | 00:30 | 12x (Stage D) |
| Extension | 72 | 00:30 | 12x (Stage D) |
| Final Extension | 72 | 07:00 | 1x |
| Hold | 4 | Infinity | 1x |

**Supplementary Table 7. Effective killing concentrations per cell line**

| Cell Line | Puromycin (µg/mL) |
|---|---|
| STS26T | 1.7 |
| 90-08 | 1.7 |
| ST-8814 | 3 |
| s462TY | 3 |
| iHSC | 1.7 |

**Supplementary Table 8. Next-Generation-Sequencing primers.**

| Name | Type | Sequence |
|---|---|---|
| P7_KERMIT_1 | i7 | CAAGCAGAAGACGGCATACGAGATTCGCCTTAGTGACTGGAGTTCAGACGTGTGCTCTTCCGATCTTCTACTATTCTTTCCCCTGCACTGT |
| P7_KERMIT_2 | i7 | CAAGCAGAAGACGGCATACGAGATCTAGTACGGTGACTGGAGTTCAGACGTGTGCTCTTCCGATCTTCTACTATTCTTTCCCCTGCACTGT |
| P7_KERMIT_3 | i7 | CAAGCAGAAGACGGCATACGAGATGCTCAGGAGTGACTGGAGTTCAGACGTGTGCTCTTCCGATCTTCTACTATTCTTTCCCCTGCACTGT |
| P7_KERMIT_4 | i7 | CAAGCAGAAGACGGCATACGAGATAGGAGTCCGTGACTGGAGTTCAGACGTGTGCTCTTCCGATCTTCTACTATTCTTTCCCCTGCACTGT |
| P7_KERMIT_5 | i7 | CAAGCAGAAGACGGCATACGAGATTTCTGCCTGTGACTGGAGTTCAGACGTGTGCTCTTCCGATCTTCTACTATTCTTTCCCCTGCACTGT |
| P7_KERMIT_6 | i7 | CAAGCAGAAGACGGCATACGAGATCATGCCTAGTGACTGGAGTTCAGACGTGTGCTCTTCCGATCTTCTACTATTCTTTCCCCTGCACTGT |
| P7_KERMIT_7 | i7 | CAAGCAGAAGACGGCATACGAGATGTAGAGAGGTGACTGGAGTTCAGACGTGTGCTCTTCCGATCTTCTACTATTCTTTCCCCTGCACTGT |
| P7_KERMIT_8 | i7 | CAAGCAGAAGACGGCATACGAGATCCTCTCTGGTGACTGGAGTTCAGACGTGTGCTCTTCCGATCTTCTACTATTCTTTCCCCTGCACTGT |
| P5_ARGON_9 | i5 | AATGATACGGCGACCACCGAGATCTACACTATAGCCTACACTCTTTCCCTACACGACGCTCTTCCGATCTTTGTGGAAAGGACGAAACACCG |
| P5_ARGON_10 | i5 | AATGATACGGCGACCACCGAGATCTACACATAGAGGCACACTCTTTCCCTACACGACGCTCTTCCGATCTTTGTGGAAAGGACGAAACACCG |
| P5_ARGON_11 | i5 | AATGATACGGCGACCACCGAGATCTACACCCTATCCTACACTCTTTCCCTACACGACGCTCTTCCGATCTTTGTGGAAAGGACGAAACACCG |
| P5_ARGON_12 | i5 | AATGATACGGCGACCACCGAGATCTACACGGCTCTGAACACTCTTTCCCTACACGACGCTCTTCCGATCTTTGTGGAAAGGACGAAACACCG |
| P5_ARGON_13 | i5 | AATGATACGGCGACCACCGAGATCTACACAGGCGAAGACACTCTTTCCCTACACGACGCTCTTCCGATCTTTGTGGAAAGGACGAAACACCG |
| P5_ARGON_14 | i5 | AATGATACGGCGACCACCGAGATCTACACTAATCTTAACACTCTTTCCCTACACGACGCTCTTCCGATCTTTGTGGAAAGGACGAAACACCG |
| P5_ARGON_15 | i5 | AATGATACGGCGACCACCGAGATCTACACCAGGACGTACACTCTTTCCCTACACGACGCTCTTCCGATCTTTGTGGAAAGGACGAAACACCG |
| P5_ARGON_16 | i5 | AATGATACGGCGACCACCGAGATCTACACGTACTGACACACTCTTTCCCTACACGACGCTCTTCCGATCTTTGTGGAAAGGACGAAACACCG |

**Supplementary Table 9. Programme gRNA cassette region PCR amplification.**

| Stage | Temperature (°C) | Time | Number of Cycles |
|---|---|---|---|
| Initial Denaturation | 98 | 03:00 | 1x |
| Denaturation | 98 | 00:15 | 5x (Stage B) |
| Annealing | 65 | 00:30 | 5x (Stage B) |
| Extension | 72 | 00:30 | 5x (Stage B) |
| Denaturation | 98 | 00:15 | 8x (Stage C) |
| Annealing | 63 | 00:30 | 8x (Stage C) |
| Extension | 72 | 00:30 | 8x (Stage C) |
| Denaturation | 98 | 00:15 | 12x (Stage D) |
| Annealing | 61 | 00:30 | 12x (Stage D) |
| Extension | 72 | 00:30 | 12x (Stage D) |
| Final Extension | 72 | 07:00 | 1x |
| Hold | 4 | Infinity | 1x |

**Supplementary Table 10. External datasets used for this study.**

| Dataset | Source |
|---|---|
| Common Essential Genes | [25] |
| DepMap gRNA Count Matrix | DepMap; [37] |
| Common non-essential Genes | DepMap; [38] |
| Screen Gene Effect scores | DepMap; [37] |
