## Supplementary Figures for "Guide RNA library representation results in gene essentiality prediction bias in genome-wide CRISPR screens"

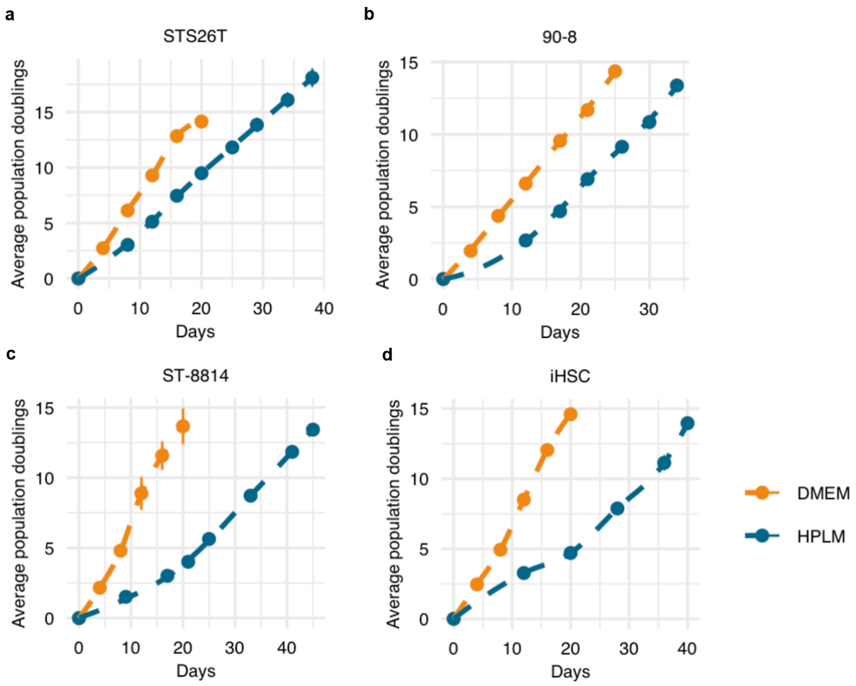


**Supplementary Figure 1. Growth curves of target cell lines in DMEM and HPLM.**

**a-d.** Growth curves of target cell lines STS26T (**a**.), 90-8 (**b**.), ST-8814 (**c**.) and iHSC (**d**.) with average population doublings (PDs) over time. Data points and dashed lines indicate PD measurements and Locally Estimated Scatterplot Smoothing (LOESS) of cells grown in DMEM (orange) and HPLM (blue). Each dot represents two biological replicates with vertical lines representing standard deviation.

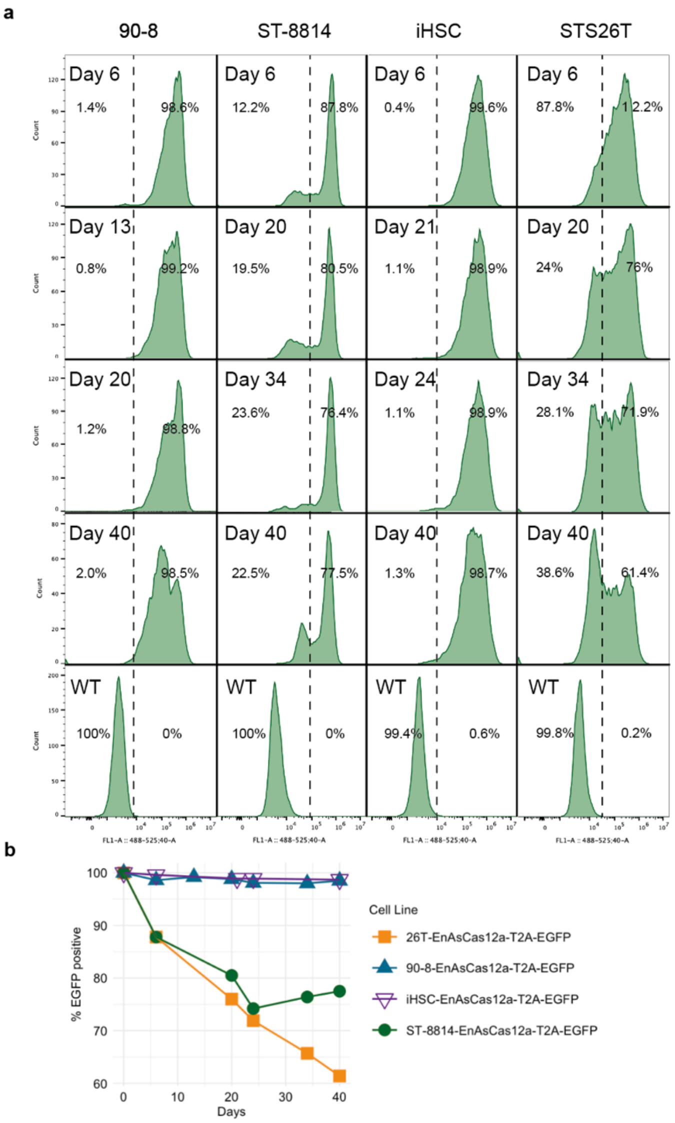


**Supplementary Figure 2. Growth curves of target cell lines in DMEM and HPLM.**

**a**. Histograms of flow cytometry measuring EGFP in 90-8, ST-8814, iHSC and STS26T cells complemented with enAsCas12a-T2A-EGFP using lentiviral transgene delivery. Cells were monitored for up to 40 days post transduction. EGFP positivity was gated stringently well above negative control populations (WT) to be able to separate true positives from pseudo positives, especially for ST-8814 and STS26T cells. (**b**) Line plot with time series data as depicted showing the percentage of EGFP positive cells based on the gating strategy as depicted in (a) over time for 90-8, ST-8814, iHSC and STS26T cells complemented with enAsCas12a-T2A-EGFP. Each data point represents one biological replicate.

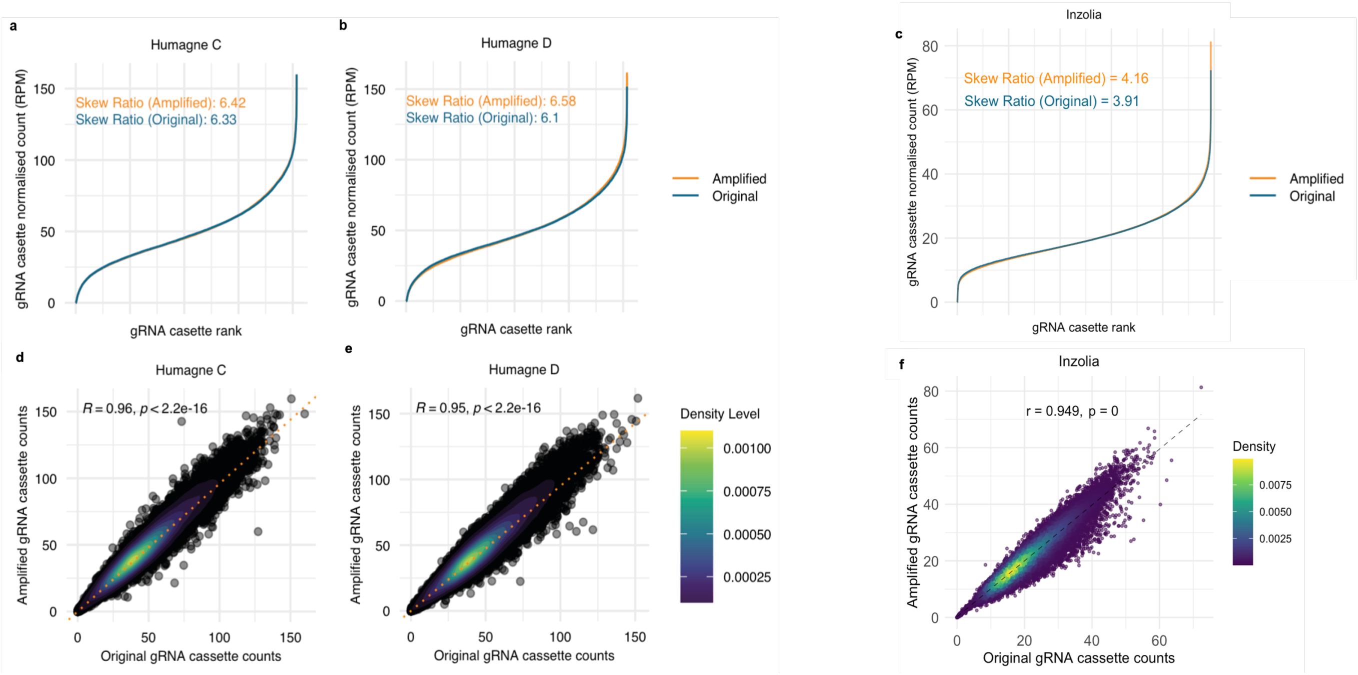


**Supplementary Figure** Error! No text of specified style in document.. **CRISPR library amplification shows minimal skewing of gRNA cassette distribution.**

**a**. gRNA cassette distribution with normalised counts per gRNA cassette for original and amplified Humagne C CRISPR libraries. The skew ratio indicates the ratio between mean normalised counts of the top 10% gRNA cassettes divided by the mean normalised counts of the bottom 10% gRNA cassettes. **b.** Same as in (a.) but for Humagne D. **c.** Same as in (a.) but for Inzolia. d. Dot and density plot of the relationship between the normalised counts of original and amplified gRNA cassettes. The spearman's rho correlation value and p-value are displayed on the plot. The Spearman's rho correlation coefficient is also depicted by an orange dashed line. **e.** Same as (d.) but for Humagne D. **f.** Same as (d.) but for Inzolia.

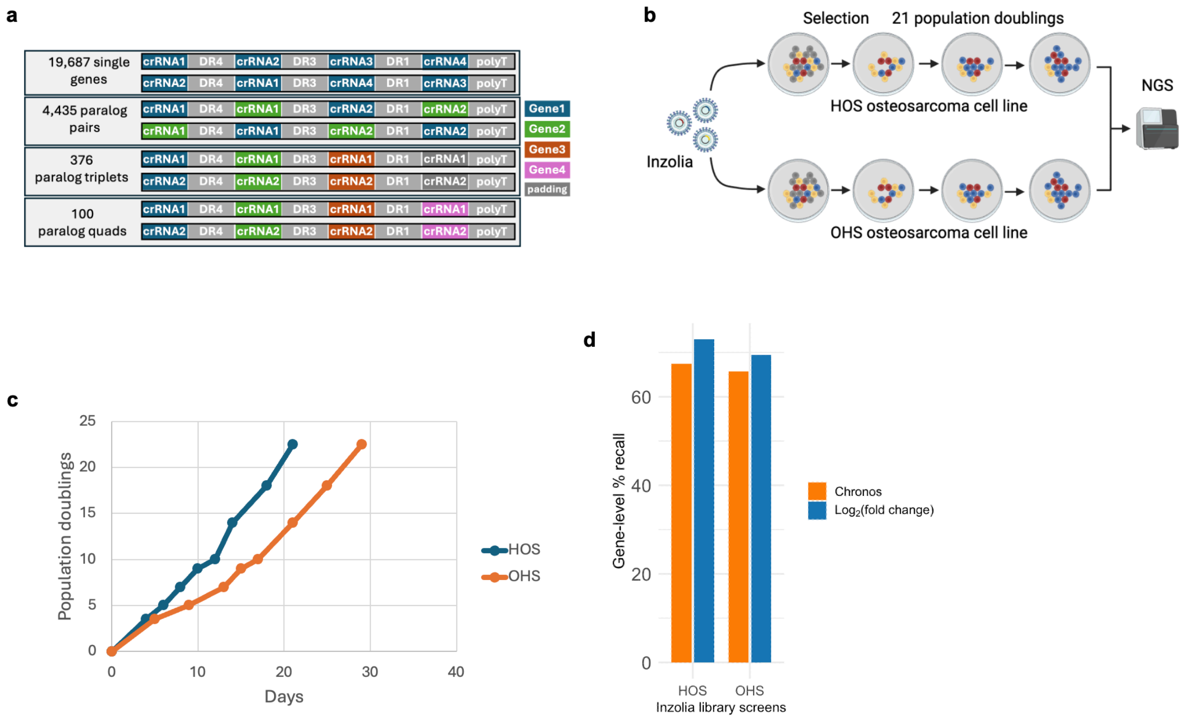


**Supplementary Figure 4. Inzolia-based CRISPR screen workflow and quality control.**

**a.** Overview of the Informa library design, which includes crRNAs for 19,687 single-gene guides, 4,435 paralog-pair guides, 376 triplet guides, and 100 quadruplet guides[32]. **b.** CRISPR screen workflow for enAsCas12a Inzolia all-in-one library for osteosarcoma cell lines HOS and OHS. **c.** Average population doublings (PDs) over time for HOS and OHS. **d.** Essential gene recall (Chronos and $\log_2$ fold change) for HOS and OHS.

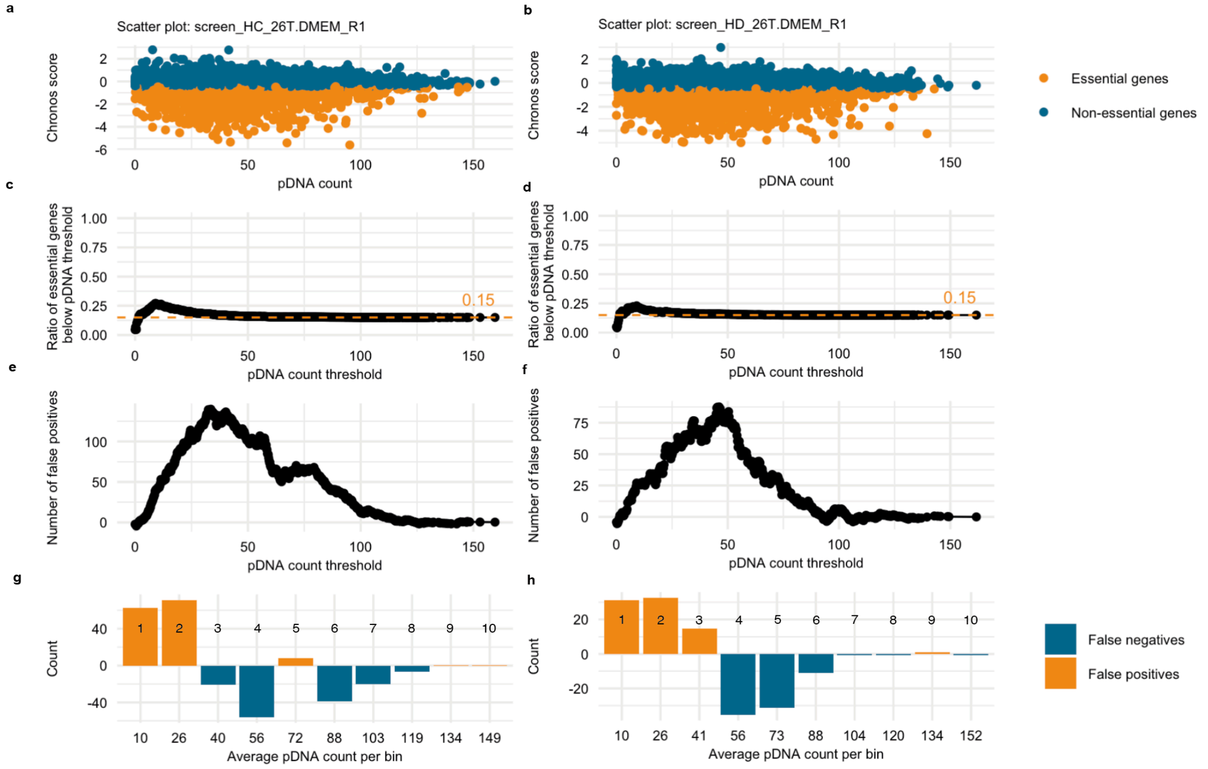


**Supplementary Figure 5. Estimation of false positive and negative results coming from pDNA representation bias. a.** Dot plot of the relationship between Chronos scores and pDNA counts in 26T, DMEM, replicate 1, Humagne C CRISPR screen. The bottom 15% genes based on Chronos score are highlighted in orange. **b.** Same as in (a.) but for the equivalent Humagne D screen. **c.** Dot plot depicting the ratio of found essential genes and the total number of genes below each possible pDNA threshold for the 26T, DMEM, replicate 1, Humagne C CRISPR screen. The dashed orange line indicates the predicted 15% essential genes threshold if essential genes were equally distributed across pDNA representation **d.** Same as in c but for the equivalent Humagne D screen. **e.** Dot plot showing the predicted number of false positives below each possible pDNA threshold for the 26T, DMEM, replicate 1, Humagne C CRISPR screen. **f.** Same as in (e.) but for the equivalent Humagne D screen. **g.** Bar graph of the estimated number of false positive and negative essential genes per pDNA bin for the 26T, DMEM, replicate 1, Humagne C CRISPR screen, with the x axis showing the average number of pDNA counts associated with each bin and the number on each bar indicating the bin number. **h.** same as (g.) but for the equivalent Humagne D screen.

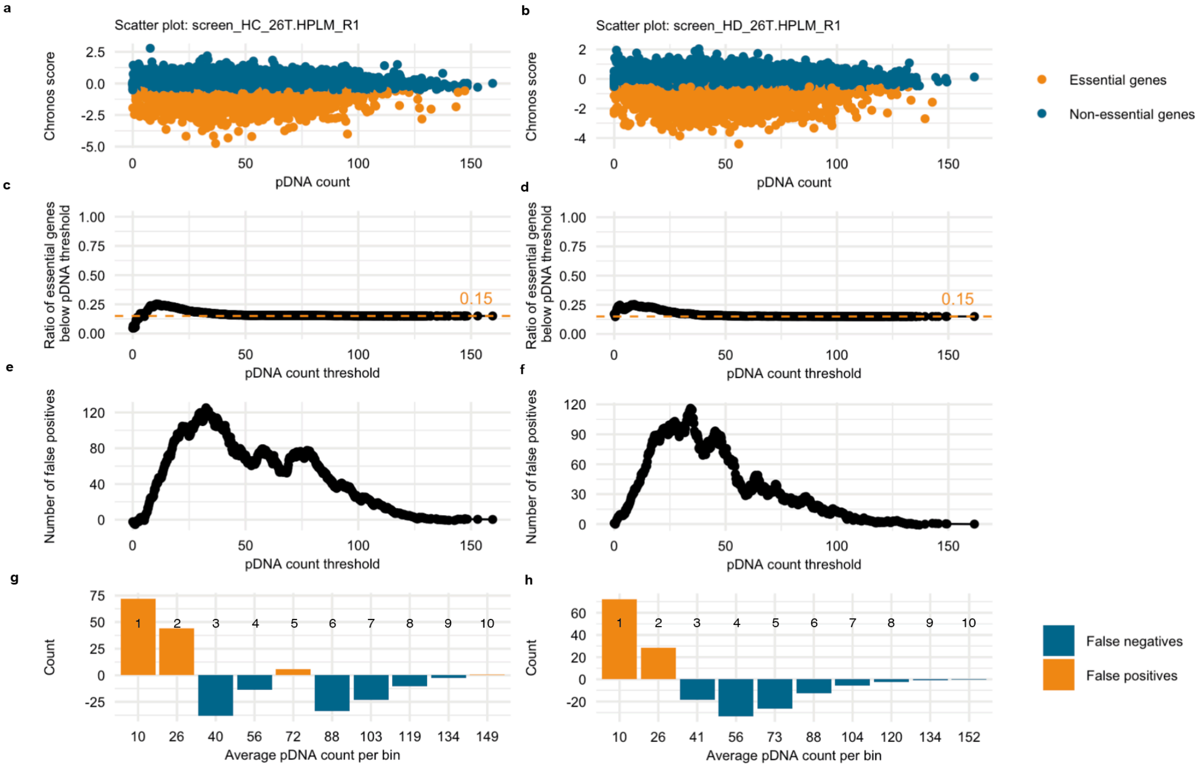


**Supplementary Figure 6. Estimation of false positive and negative results coming from pDNA representation bias. a-h**: same as Supplementary figure 4, but for the STS26T, HPLM, replicate 1 screen.

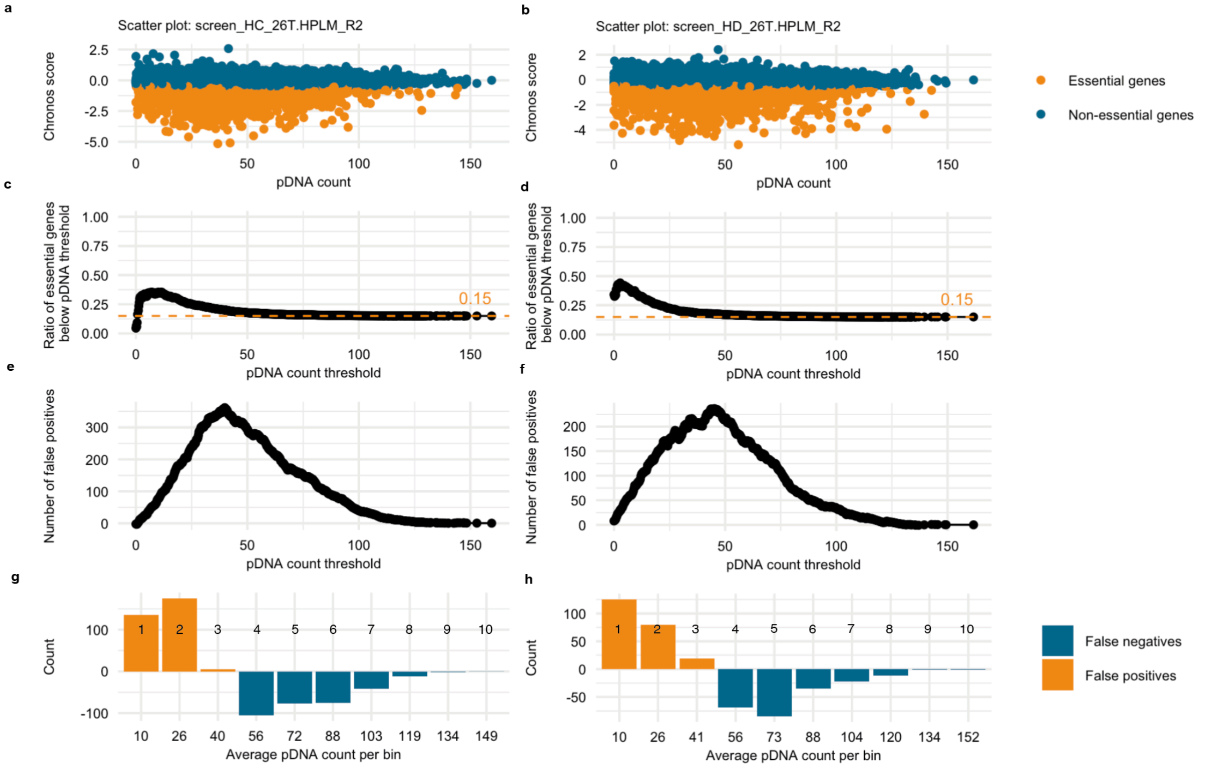


**Supplementary Figure 7. Estimation of false positive and negative results coming from pDNA representation bias. a-h**: same as Supplementary figure 4, but for the STS26T, HPLM, replicate 2 screen.

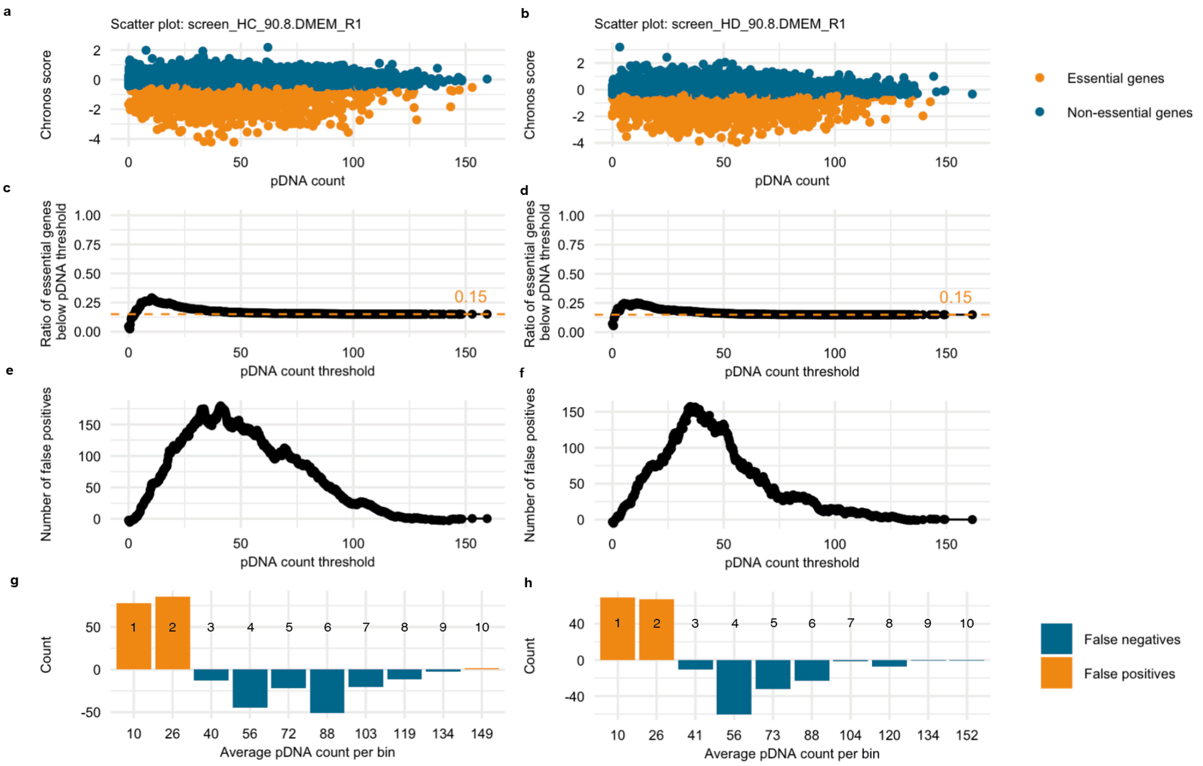


**Supplementary Figure 8. Estimation of false positive and negative results coming from pDNA representation bias. a-h**: same as Supplementary figure 4, but for the 90-8, DMEM, replicate 1 screen.

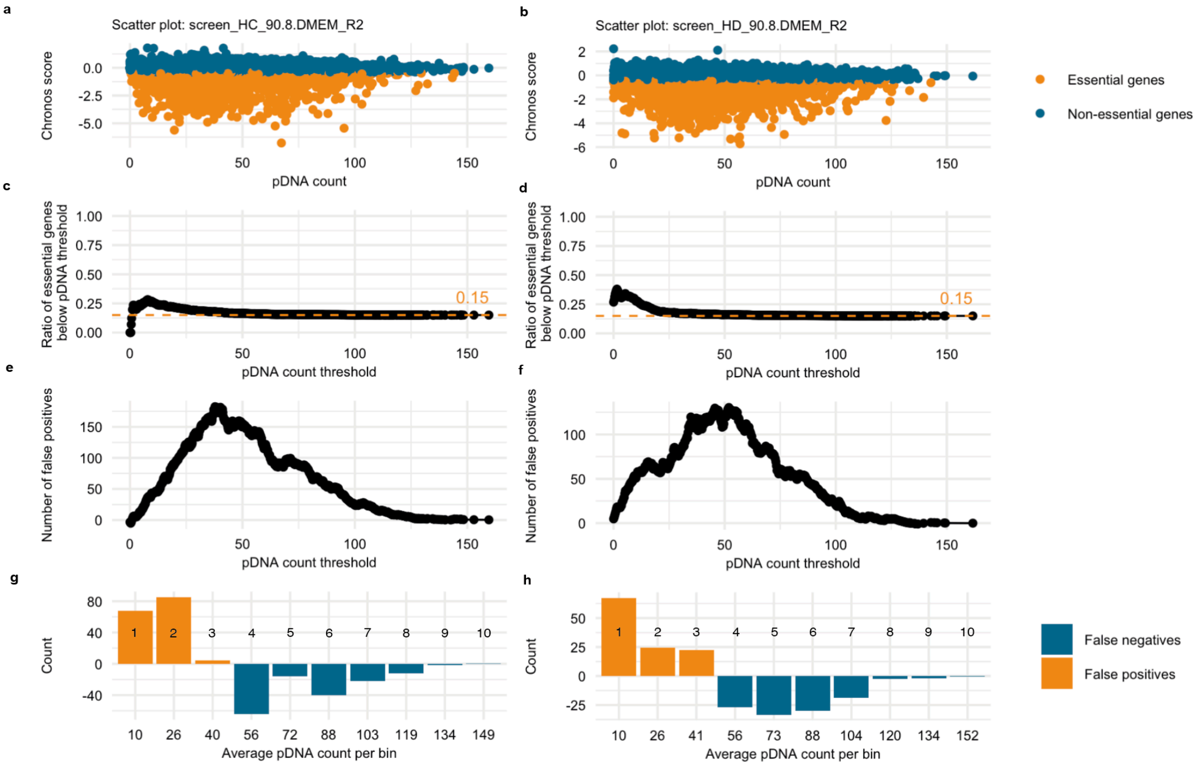


**Supplementary Figure 9. Estimation of false positive and negative results coming from pDNA representation bias. a-h**: same as Supplementary figure 4, but for the 90-8, DMEM, replicate 2 screen.

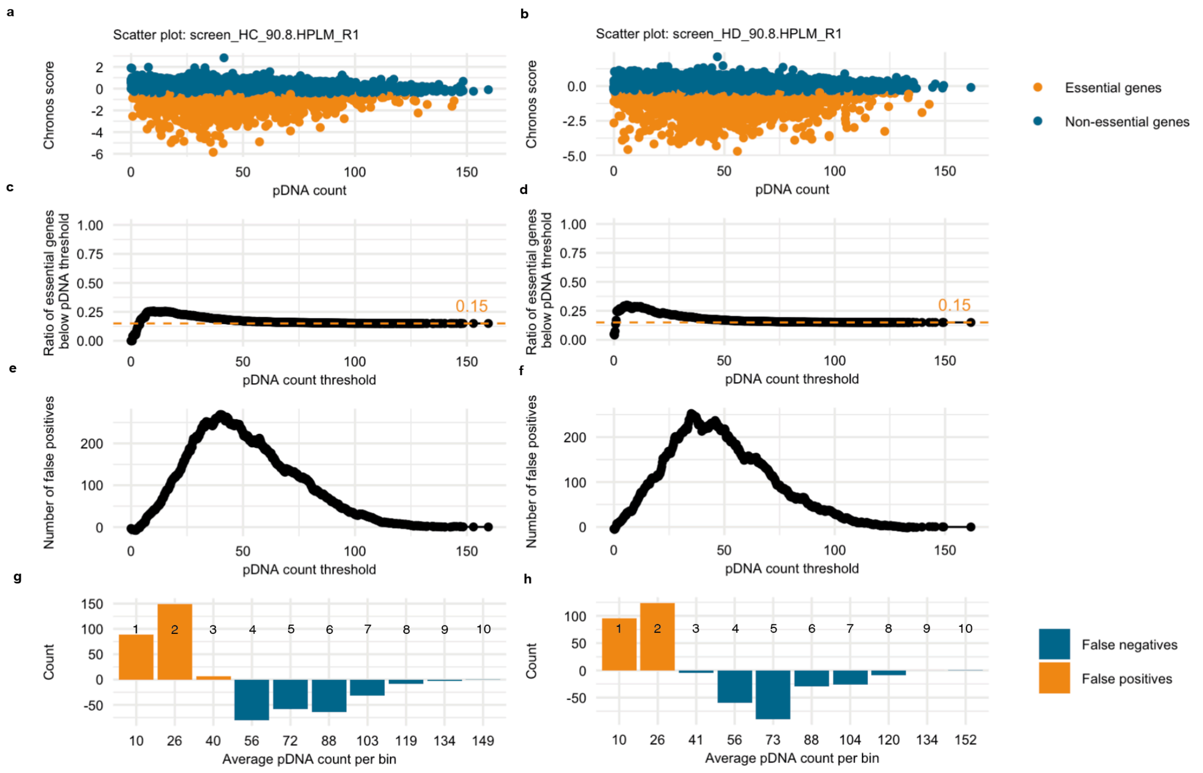


**Supplementary Figure 10. Estimation of false positive and negative results coming from pDNA representation bias. a-h**: same as Supplementary figure 4, but for the 90-8, HPLM, replicate 1 screen.

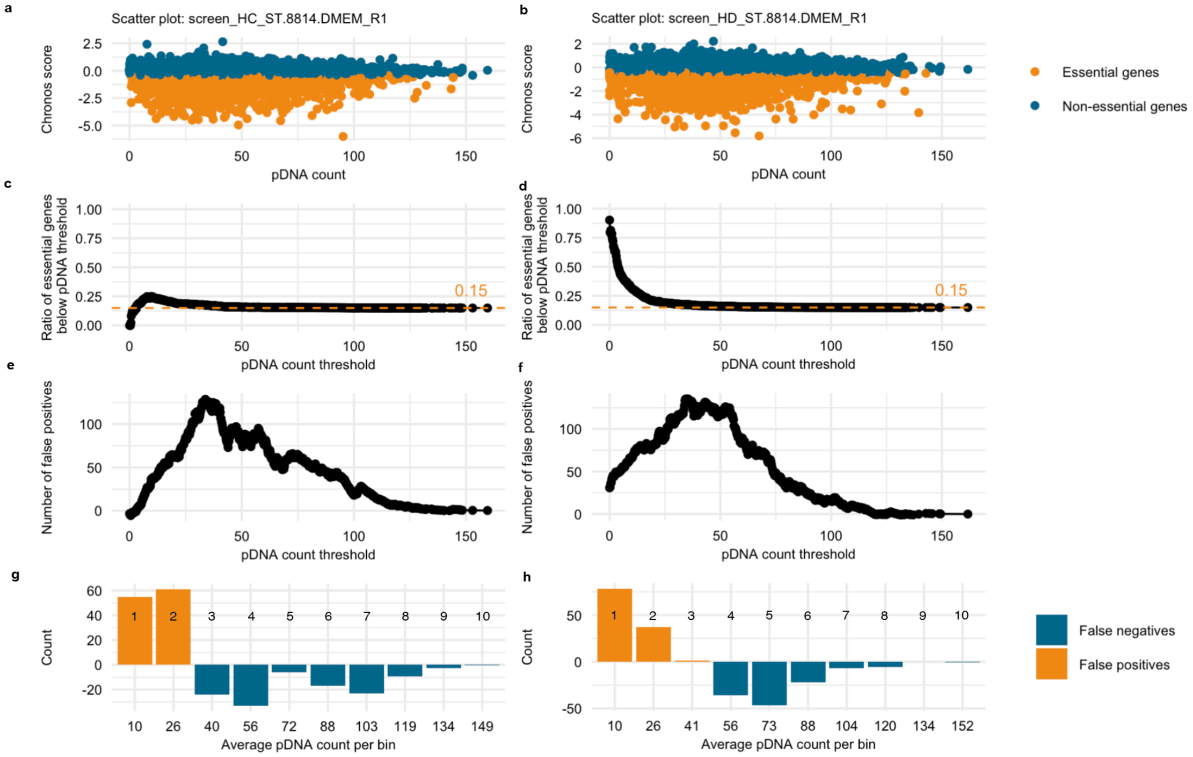


**Supplementary Figure 11. Estimation of false positive and negative results coming from pDNA representation bias. a-h**: same as Supplementary figure 4, but for the ST-8814, DMEM, replicate 1 screen.

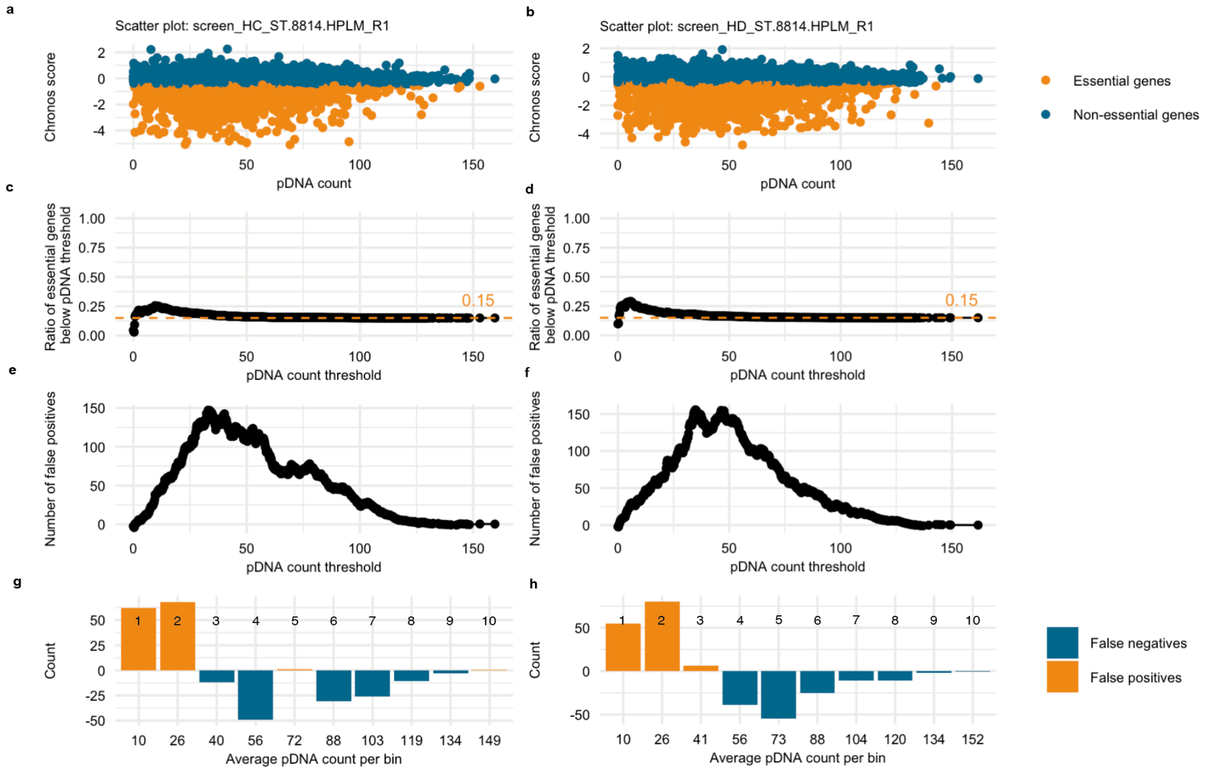


**Supplementary Figure 12. Estimation of false positive and negative results coming from pDNA representation bias. a-h**: same as Supplementary figure 4, but for the ST-8814, HPLM, replicate 1 screen.

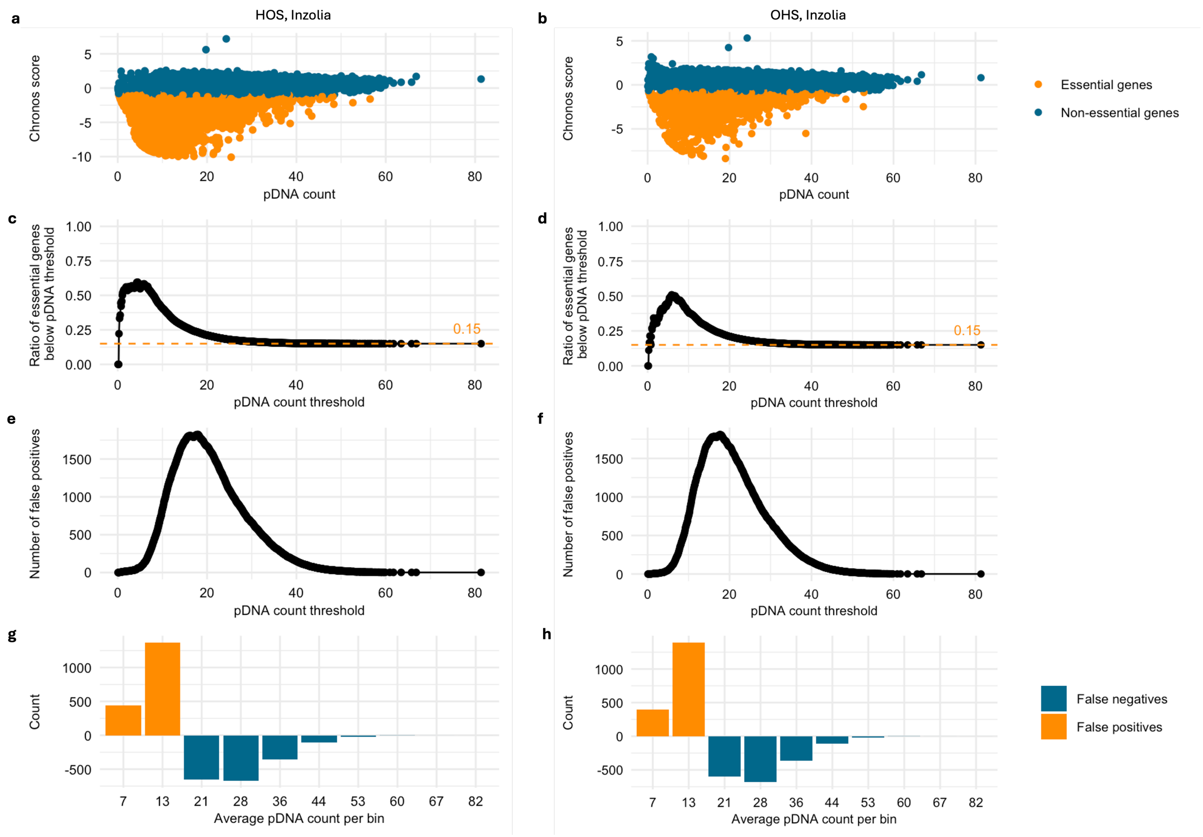


**Supplementary Figure 13. Estimation of false positive and negative results coming from pDNA representation bias. a-h**: same as Supplementary figure 4, but for Inzolia-based CRISPR screens in HOS and OHS cell lines.
