## Supplementary material for "Guide RNA library representation results in gene essentiality prediction bias in genome-wide CRISPR screens": Cover Letter

7th September 2025

The Editors

**Guide RNA library representation results in gene essentiality prediction bias in genome-wide CRISPR screens.**

Paul Metz[1∫], Sofia Alves-Vasconcelos[1], Richard Wallbank[1], Joey Riepsaame[1,2] and A. Bassim Hassan[1,¶].

[1]Oxford Molecular Pathology Institute, Sir William Dunn School of Pathology, University of Oxford, South Parks Road, Oxford, OX1 3RE, United Kingdom.
[2]Genome Engineering Oxford, Sir William Dunn School of Pathology, University of Oxford, South Parks Road, Oxford, OX1 3RE, United Kingdom.

Dear Sir/Madam,

We would be most grateful if you would consider the above manuscript for publication in your Journal. We justify our submission because we believe this work to be of general interest to a wide range of research fields that utilise genome wide CRISPR perturbation screens, including developmental biology, cancer biology and translational biology.

The background to our laboratory is signalling in development and cancer, where we have previously focused on the IGF2/Mannose 6-phosphate receptor interactions with the IGF2 growth factor ligand using structural biology (e.g. EMBOJ 2002&2007, Science 2012, PNAS 2016, Structure 2020). We have also utilised conditional mouse genetics to evaluate *Tp53*, *Cdh1*, *Igf2r* and *Smad4* function (e.g. EMBO Mol Med 2012, Oncotarget 2016, Sci Rep 2019, Sci Rep 2025).

Genome wide CRISPR-based perturbation screens are powerful discovery tools enabling the identification of novel gene dependencies through either gain or loss of function approaches. While genome wide guide RNA libraries have advantages when using enAsCas12a, such as multiplex single guides per gene, they may be subject to similar confounding factors that can affect the interpretation of large genome-wide datasets. Here, we examine the impact of these variables in over twenty enASCas12a gRNA perturbation screens that we have performed using Humagne C, Humagne D and Inzolia libraries in human cells. We demonstrate that the choice of CRISPR library is often the most significant factor that influences genetic perturbation results, outweighing other variables such as either target cell lines or culture media conditions. A major contributor to this effect is guide RNAs (gRNAs) representation bias within a given CRISPR library, where lower gRNA representation, or plasmid DNA representation (pDNA), can lead to variable and more pronounced gene effect scores using either log fold change or Chronos analysis. The observations also extend to DepMap Cas9 screen datasets in DepMap as well as enCas12a. These effects may be mitigated by using either multiple gRNA constructs per gene, by optimisation of CRISPR library production processes or by targeting with multiple independent gRNA libraries. Importantly, we also consider gRNA representation bias during CRISPR screen hit prioritisation. CRISPR library gRNA representation bias remains a major challenge in the interpretation of gene essentiality in perturbation screens.

Importantly, we estimated the number of false positives and negative essential genes based on the expected number of essential genes and the observed number of essential genes in different gRNA representation categories. In line with our findings, the number of essential genes, as defined by the bottom 15% scoring genes, is higher amongst genes associated with gRNAs that have a relatively low initial representation. By estimating the number of false positive and negative essential genes per gRNA representation bin, we predicted that the total number of wrongly classified (false positive or negative) genes in screen data from this study varies between 6 and 23%.

Aside from PCR cycle optimisation and PCR primer sequence bias, reducing gRNA representation bias in modest scale screens would include either the use of larger scale screens to compensate for low gRNA representation or to utilise independently generated guide libraries, that fundamentally differ in pDNA representation, combined with elimination of poorly represented gRNAs. For a given screen coverage and cell number, these would still have cost-efficiency implications and result in either multiple library screens per experiment or even pooled but separately generated gRNA libraries with high cell numbers.

We thank you for your consideration in reviewing the manuscript and we would be delighted to address any further questions concerning the work. We have suggested manuscript reviewer's names below.

Yours sincerely.

Professor Bass Hassan
On behalf of the authors,
University of Oxford

**Suggested Reviewers to invite**
Claudia Feng, Genome Biology Associate Editor
Professor Eric Chow, Laboratory for Genomics Research, USCF.
Professor Chris Lord, Institute of Cancer Research, London.

**Suggested Reviewers to not to invite**
Josh Dempster, The Broad Institute, Harvard University
John Doench, The Broad Institute, Harvard University
